# More than *mcr*: Canonical Plasmid- and Transposon-Encoded Mobilized Colistin Resistance (*mcr*) Genes Represent a Subset of Phosphoethanolamine Transferases

**DOI:** 10.1101/2022.10.03.510586

**Authors:** Ahmed Gaballa, Martin Wiedmann, Laura M. Carroll

**Author notes:** **Correspondence:** Laura M. Carroll.

## Abstract

Mobilized colistin resistance genes (*mcr*) may confer resistance to colistin, a last-resort, critically important antimicrobial for human health. *mcr* can often be transmitted horizontally (e.g., via mobile genetic elements); however, *mcr* encode phosphoethanolamine transferases (PET) closely related to chromosomally encoded, intrinsic lipid modification enzymes (e.g., EptA, EptB, CptA). To explore the genetic diversity of *mcr* within the context of intrinsic lipid modification PET, we identified 9,836 non-redundant protein accession numbers associated with *mcr*-like genes, representing a total of 69,814 *mcr*-like genes present across 256 bacterial genera. We subsequently identified 125 unique, putative novel *mcr*-like genes encoded on the same contig as a plasmid replicon and other antimicrobial resistance genes. Sequence similarity and a maximum likelihood phylogeny of *mcr*, putative novel *mcr*-like genes, and intrinsic lipid modification PET-encoding genes indicated that sequence similarity is insufficient to discriminate between genes involved in colistin resistance and genes encoding intrinsic lipid modification PET. A mixed-effect model of evolution (MEME) indicated that site- and branch-specific diversifying positive selection might have played a role in the evolution of subvariants within the *mcr-2* and *mcr-9* families. MEME suggested that positive selection played a role in the diversification of several residues in structurally important regions, including (i) a bridging region that connects the membrane-bound and catalytic periplasmic domains, and (ii) a periplasmic loop juxtaposing the substrate entry tunnel. These residues were found to be differentially conserved in different *mcr* families and thus may play a role in *mcr* subvariant phenotypic diversity. Moreover, we found that *eptA* and *mcr* are localized within different genomic contexts. Canonical *eptA* are typically chromosomally encoded in an operon with a two-component regulatory system or adjacent to a TetR-type regulator. In contrast, *mcr* are encoded as single-gene operons or adjacent to *pap2* and *dgkA*, which encode a PAP2 family lipid A phosphatase and diacylglycerol kinase, respectively. Our data suggest that *eptA* can give rise to “colistin resistance genes” through various mechanisms, including selection and diversification of the genomic context, regulatory pathways, and mobilization. These mechanisms likely altered gene expression levels and enzyme activity, allowing bona fide *eptA* to evolve to function in colistin resistance.

## 1 Introduction

Colistin is a polycationic peptide, which serves as a last-resort antimicrobial used to treat infections caused by multidrug-resistant (MDR), extensively drug-resistant (XDR), and pan-drug-resistant (PDR) Gram-negative bacteria (El-Sayed Ahmed et al., 2020; Binsker et al., 2022). Consequently, colistin has been designated by the World Health Organization (WHO) as a highest priority critically important antimicrobial for human medicine (World Health Organization, 2019). Due to the critical importance of colistin as an antibiotic of last resort, colistin resistance among Gram-negative pathogens represents an increasingly dire global public health threat (El-Sayed Ahmed et al., 2020; World Health Organization, 2021; Binsker et al., 2022).

The binding of colistin to bacterial cells is initiated by an electrostatic attraction between the colistin cationic head group and the anionic phosphate group on the lipid A of the bacterial lipopolysaccharide (LPS), which displaces Ca^2+^ and Mg^2+^ ions (Olaitan et al., 2014; El-Sayed Ahmed et al., 2020). Subsequently, colistin’s hydrophobic tail integration in the lipid bilayers leads to cell membrane disruption and cell death by targeting cytoplasmic membrane LPS (Trimble et al., 2016; Sabnis et al., 2021). Multiple colistin resistance mechanisms have been reported, including (i) modification of lipid A (Guo et al., 1997; Gunn et al., 1998); (ii) expression of a multi-drug efflux system in *Pseudomonas aeruginosa* (AbuOun et al., 2017); (iii) the complete loss of LPS in *Acinetobacter baumannii* (Moffatt et al., 2010); (iv) overproduction of capsular polysaccharide in *Klebsiella pneumoniae* (Campos et al., 2004); and (v) enzymatic degradation of colistin via colistin-degrading proteases in *Stenotrophomonas maltophilia* (Ito-Kagawa and Koyama, 1980; Lee et al., 2022). The main mechanism of bacterial colistin resistance is the modification of lipid A via the addition of a cationic group, such as phosphoethanolamine (pEtN), 4-amino-4-deoxy-L-arabinose, and glycine to lipid A (Trimble et al., 2016), which reduces the overall membrane negative charge and subsequently decreases colistin binding affinity to the cell (Olaitan et al., 2014; El-Sayed Ahmed et al., 2020). While it has been shown that lipid A is essential for growth in most bacterial species, lipid A modifications are dispensable for cell survival under laboratory growth conditions (Trent, 2004; Raetz et al., 2007; Parsons and Rock, 2013). However, lipid A modification plays a significant role in bacterial adaptation to different stress conditions, including mild acid stress, change in oxygen level, increase in growth temperature, pH change, osmotic stress, and the presence of cationic antimicrobial compounds (Trent, 2004; Raetz et al., 2007; Gunn, 2008; Anandan and Vrielink, 2020; Troudi et al., 2021).

Some Gram-negative bacteria (e.g., *Neisseria, Serratia, Brucella, Burkholderia* spp.) showcase intrinsic resistance to colistin; others, including pathogenic Enterobacteraceae, can acquire resistance via numerous mechanisms, including chromosomal mutations (e.g., those that modify the bacterial cell surface) and/or mobilized colistin resistance genes (*mcr*) (El-Sayed Ahmed et al., 2020; World Health Organization, 2021; Binsker et al., 2022). *mcr* represent a particularly notable and concerning mechanism of colistin resistance, as they are predominantly plasmid-borne and/or associated with insertion sequences and can facilitate the acquisition and rapid dissemination of colistin resistance (Kieffer et al., 2017; Trebosc et al., 2019; El-Sayed Ahmed et al., 2020). *mcr* encodes an inner membrane-anchored protein that belongs to a complex group of phosphoethanolamine transferase (PET) enzymes (Gao et al., 2016), which adds phosphoethanolamine (pEtN) to different moieties of lipid A (Cox et al., 2003; Reynolds et al., 2005; Tamayo et al., 2005; Raetz et al., 2007; Parsons and Rock, 2013; Anandan et al., 2017). Gram-negative bacterial species express one or more housekeeping PET, which function in intrinsic lipid modification, including EptA, EptB, and CptA (Anandan and Vrielink, 2020). We will collectively refer to *eptA, eptB*, and *cptA* genes as *ipet* (**<u>i</u>**ntrinsic lipid modification **<u>PET</u>**) and their product as i-PET. Lipid A modifying i-PET enzymes phylogenetically cluster into three distinct groups that correlate with the enzyme specificity to the pEtN-acceptor substrate site on lipid A. For example, (i) EptA adds pEtN to the N-acetylglucosamine moiety of lipid A, (ii) EptB adds pEtN to the 3-deoxy-d-manno-octulosonic acid (KDO) of the LPS core region, and (iii) CptA/EptC adds pEtN to the second heptose of the LPS core region (Harper et al., 2017; Anandan and Vrielink, 2020).

Interestingly, *mcr* share a high degree of sequence and structural similarity with i-PET enzymes present in many Gram-negative bacteria (Bakovic et al., 2007; Anandan and Vrielink, 2020; Samantha and Vrielink, 2020), including pathogenic members of Enterobacteriaceae (El-Sayed Ahmed et al., 2020; Gogry et al., 2021). PET protein structure includes two discretely folded domains connected by a bridging helix and extended loop, an N-terminal transmembrane domain, and a C-terminal periplasmic soluble catalytic domain (Anandan et al., 2017; Anandan and Vrielink, 2020). Moreover, PET are metalloenzymes that share a conserved zinc-binding site tetrahedrally coordinated by the side chains of conserved residues Glu^246^, Thr^285^ His^466,^ and Asp^465^ (numbers corresponding to the MCR-1 sequence), with the Thr^285^ residue acting as the catalytic nucleophile for the pEtN transfer (Stojanoski et al., 2016; Anandan et al., 2017). While multiple conserved or partially conserved amino acid (AA) residues were suggested to be involved in pEtN binding (Sun et al., 2017; Dortet et al., 2018; Stogios et al., 2018; Carroll et al., 2019; Garcia-Menino et al., 2020), the lipid-binding pocket remains poorly defined (Carroll et al., 2019).

As of April 2022, >95 subvariants belonging to 10 major *mcr* families (termed *mcr-1* to *-10*, which encode MCR-1 to -10, respectively) have been identified across various Gram-negative bacterial genera (Liu et al., 2016; Xavier et al., 2016; AbuOun et al., 2017; Borowiak et al., 2017; Carattoli et al., 2017; Yin et al., 2017; Wang et al., 2018b; Yang et al., 2018; Carroll et al., 2019; Wang et al., 2020). While MCR subvariants share a conserved overall protein structure (Stojanoski et al., 2016; Sun et al., 2017; Xu et al., 2018a; Carroll et al., 2019; Son et al., 2019), MCR-encoding genes share varied degrees of sequence similarities (Li et al., 2018; Carroll et al., 2019; Ramaloko and Osei Sekyere, 2022). Moreover, it has been shown that different *mcr* subvariants vary in terms of the colistin resistance levels they confer (Nang et al., 2019; El-Sayed Ahmed et al., 2020). However, the molecular bases of *mcr* genetic and phenotypic heterogeneities are poorly understood. Furthermore, it has been suggested that *mcr* evolved from an *eptA* chromosomal gene copy via mobilization, and that *Moraxella* spp. are a potential source for MCR-like colistin resistance determinants (Kieffer et al., 2017). Indeed, several lines of evidence support the notion that *mcr* evolved from *eptA* through mobilization, including: (i) MCR and EptA share a conserved overall protein structure and acceptor substrate specificity, (ii) ISApl1-dependent mobilization of *mcr* has been demonstrated (AbuOun et al., 2017; Kieffer et al., 2017), and (iii) insertion of ISApl1 upstream of *eptA* increased *eptA* expression and led to an increased level of colistin resistance in XDR *Acinetobacter baumannii* isolates (Trebosc et al., 2019). However, the exact ancestor of *mcr* and the molecular mechanism of evolution of intrinsic lipid modification *eptA* to mobilized colistin-resistant determinant remains unknown.

Here, we aim to provide further insight into the evolutionary relationships between *mcr* subvariants, all within the context of i-PET. We show that some i-PET (e.g., *eptA*) may differ from canonical *mcr* determinants in terms of their genomic context, and we identify differentially conserved residues, which may play a role in the levels of colistin resistance conferred by different *mcr* subvariants. Finally, using a large number (>69,000) of *mcr*-like genes extracted from publicly available bacterial genomes, we identify several putative, novel *mcr*-like genes, which are likley co-harbored on plasmids with other antimicrobial resistance (AMR) genes. Overall, the results presented here provide insight into the evolution and emergenece of *mcr* subvariants in bacteria.

## 2. Materials and Methods

### 2.1 Acquisition of *mcr* and *mcr*-like amino acid sequences

Amino acid (AA) sequences of nine published *mcr* homologs (*mcr-1* to *-9*; Supplementary Table S1) were queried against the National Center for Biotechnology Information (NCBI) non-redundant (nr) protein database using the protein BLAST (BLASTP) webserver (https://blast.ncbi.nlm.nih.gov/Blast.cgi?PAGE=Proteins) (Johnson et al., 2008), using the default settings for all parameters except max target sequences, which was raised to 5,000 (accessed April 23-24, 2019). Only hits corresponding to proteins with high query coverage (i.e., ≥90% of the length of the original *mcr* query) were maintained (*n* = 41,270 out of 45,001 total hits). From this, the union of all nr protein accession numbers was taken, yielding a total of 9,866 nr protein accession numbers associated with *mcr* and *mcr*-like genes. The AA sequences of all 9,866 *mcr* and *mcr*-like genes, as well as all associated NCBI Identical Protein Group (IPG) accession numbers, were downloaded using the rentrez package version 1.2.1 (Winter, 2017) in R version 3.5.3 (R Core Team, 2019). Finally, to remove extreme outliers, the median absolute deviation of the lengths of the AA sequences of all *mcr* and *mcr*-like genes was calculated, and AA sequences of *mcr*-like genes falling outside the range of sequence lengths encompassed by 15 times the median absolute deviation were removed, yielding a total of 9,836 AA sequences used in subsequent steps.

### 2.2 Acquisition and characterization of genomes harboring *mcr*-like genes

NCBI RefSeq Assembly accession numbers for all genomes associated with ≥1 *mcr*-like gene in NCBI’s IPG database were acquired via rentrez (see section “Acquisition of *mcr* and *mcr*-like amino acid sequences” above). Latest assembly versions for all RefSeq genomes were then downloaded via NCBI’s FTP site (*n* = 59,129 total genomes; accessed May 19, 2020). To confirm that ≥1 *mcr*-like gene could be detected in each genome, the BLASTX algorithm implemented in DIAMOND version 0.9.13.114 (Buchfink et al., 2015) was used to perform a translated search of each genome against AA sequences of all 9,836 *mcr*-like genes identified as described above (see section “Acquisition of *mcr* and *mcr*-like amino acid sequences” above), plus the AA sequences of all 53 known *mcr* subvariants representing *mcr-1* to *-9* available in ResFinder (https://cge.cbs.dtu.dk/services/ResFinder/) at the time (*n* = 9,889 total *mcr* and *mcr-*like genes; accessed April 24, 2019). The following DIAMOND BLASTX parameters were used to confirm *mcr* and *mcr*-like gene presence: a maximum E-value threshold of 1e-5 (-e 0.00001), a minimum subject coverage threshold of 90% (--subject-cover 90), a minimum percent AA identity of 90% (--id 90). This resulted in a total of 69,814 confirmed hits of *mcr* and *mcr-*like genes in 59,121 RefSeq genomes, which were used in subsequent steps (Supplementary Table S2).

To assign each genome to a species using a standardized taxonomic framework, all 59,121 genomes were assigned to a marker-gene based operational taxonomic unit (mOTU) using classify-genomes (accessed June 3, 2020; https://github.com/AlessioMilanese/classify-genomes) and version 2.5 of the mOTUs taxonomy (Milanese et al., 2019) (Supplementary Table S2). Antimicrobial resistance (AMR) genes and plasmid replicons were detected in each genome using ABRicate version 0.9.8 (https://github.com/tseemann/abricate), plus the NCBI National Database of Antibiotic Resistant Organisms (NDARO) and PlasmidFinder database, respectively (each accessed April 19, 2020) (Carattoli et al., 2014; Feldgarden et al., 2019). AMR genes and plasmid replicons were considered to be present in a genome using minimum nucleotide identity and coverage thresholds of 80% each (Supplementary Table S2). PlasFlow version 1.1.0 (Krawczyk et al., 2018) was additionally used to predict whether contigs were plasmid-associated or chromosomal in origin (using default settings; Supplementary Table S2).

### 2.3 Identification of putative novel *mcr*-like genes

Of the 69,814 *mcr* and *mcr*-like genes identified in 59,121 genomes from NCBI’s RefSeq database (see section “Acquisition and characterization of genomes harboring *mcr*-like genes” above), we identified 321 *mcr*-like genes, which were located on the same contig as (i) ≥1 plasmid replicon and (ii) ≥1 additional AMR gene, with contigs harboring previously described *mcr* subvariants excluded (see section “Acquisition and characterization of genomes harboring *mcr*-like genes” above for details regarding plasmid replicon and AMR gene detection; Supplementary Table S2). These 321 genes, which we will refer to hereafter as “putative novel *mcr*-like genes”, represented 125 unique nucleotide sequences, which were used in subsequent steps (Supplementary Table S3).

### 2.4 Acquisition of chromosomally encoded i-PET genes

Closed representative genome sequences of 147 reference species known to encode *mcr* genes were downloaded from the NCBI RefSeq Assembly database (accessed March 12, 2021). These included Enterobacteriaceae (NCBI txid543), *Proteus* (NCBI txid583), *Aeromonas* (NCBI txid642), and *Moraxella* (NCBI txid475) species (Supplementary Table S4). Assembled genomes were imported into Geneious version 2019.2.3 (Biomatters, Auckland, New Zealand), and sequences of complete closed chromosomes were extracted for subsequent analysis. AA sequences of *E. coli* str. K-12 substr. MG1655 CptA (NCBI Protein Accession WP_000556306.1), EptA (NCBI Protein Accession WP_000919792.1), and EptB (NCBI Protein Accession WP_001269197.1) were queried against the closed chromosome sequences using translated nucleotide BLAST (tBLASTN, as implemented in Geneious version 2019.2.3; Biomatters, Auckland, New Zealand). A total of 237 unique hits that showed ≥90% query coverage were selected to represent chromosomally encoded i-PET genes (Supplementary Table S5). We will collectively refer to *eptA, eptB*, and *cptA* intrinsic lipid modification PET-encoding genes as *ipet* and their products as i-PET.

### 2.5 Construction of maximum likelihood phylogeny of *mcr* and *mcr*-like genes

Maximum likelihood (ML) phylogenies were inferred from the nucleotide sequences of (i) all *mcr* subvariants available in NCBI’s NDARO (*n* = 98, accessed March 12, 2021; Supplementary Table S6), (ii) putative novel *mcr*-like genes identified in this study (*n* = 125; see section “Identification of putative novel *mcr*-like genes” above) and/or (iii) *ipet* (*n* = 237) coding sequences (see section “Acquisition of chromosomally encoded i-PET genes” above; Supplementary Table S7). Back-translated nucleotide multiple sequence alignments (NT_btn_-MSA) were constructed using MUSCLE (Edgar, 2004) with the default settings in Geneious version 2019.2.3 (Biomatters, Auckland, New Zealand). The resulting NT_btn_-MSAs were used to construct ML phylogenies with 100 bootstrap replicates via RAxML, using the GTRGAMMA substitution model and default settings (RAxML GUI version 2.0.7 and RAxML version 8.2.12) (Stamatakis, 2014). The resulting trees were visualized and edited using iTOL version 6.5 (https://itol.embl.de/) (Letunic and Bork, 2007).

### 2.6 Descriptive sequence analysis

The number of polymorphic sites, nucleotide diversity per site, average pairwise nucleotide differences per sequence, number of synonymous substitutions (S) and non-synonymous substitutions (N), and the *dN/dS* ratio for each NT_btn_-MSA (Supplementary Table S7; see section “Construction of maximum likelihood phylogeny of *mcr* and *mcr*-like genes” above) were calculated using DnaSP version 6.12.03 (Rozas et al., 2017).

### 2.7 Homologous recombination detection

The Recombination Detection Program (RDP5) was used to detect homologous recombination events withn the NT_btn_-MSA of *mcr*, putative novel *mcr*-like genes, and *ipet* (*n* = 98, 125, and 237 genes, respectively; Supplementary Table S7) (Martin et al., 2021). Seven homologous recombination detection methods (RDP, BOOTSCAN, GENECON V, MAXCHI, CHIMAERA, SISCAN, and 3Seq) were used to perform a full exploratory recombination analysis using the default settings in RDP5. Only recombination events detected by at least three methods were selected for further analysis to reduce false positives. Recombination events with similar breakpoints were merged into a single event, and the overall significance of the recombination evidence for each event was evaluated via the Pairwise Homoplasy Index (PHI) test with default settings in RDP5.

### 2.8 Positive selection analysis

The Datamonkey server (https://www.datamonkey.org/) (Weaver et al., 2018) was used to identify *mcr* subvariant residues that evolved under diversifying selection. The “hyphy cln” command within HyPhy 2.5.32(MP) (Kosakovsky Pond et al., 2020) was used to remove stop codons within the NT_btn_-MSA of the 98 known *mcr* subvariants (Supplementary Table S6). The resulting cleaned alignment was supplied as input to the command line implementation of GARD (Kosakovsky Pond et al., 2006; Weaver et al., 2018; Kosakovsky Pond et al., 2020), which was used to detect recombination breakpoints within the alignment using default settings. GARD partitioned the dataset into fragments of 1-1,204 bp and 1,205-1,779 bp.

The resulting partitioned dataset produced by GARD (with suffix .best-gard) was supplied as input to the following (both accessed January 27, 2022): (i) FUBAR (Fast, Unconstrained Bayesian AppRoximation; https://www.datamonkey.org/fubar), which uses a Bayesian approach to infer non-synonymous (*dN*) and synonymous (*dS*) substitution rates on a per-site basis and assumes constant selection pressure for each site along the entire phylogeny (Murrell et al., 2013); (ii) MEME (mixed-effect model evolution-based diversifying selection analysis; https://www.datamonkey.org/meme), which tests the hypothesis that individual sites have been subject to episodic positive or diversifying selection (Murrell et al., 2012). For both FUBAR and MEME, the universal genetic code option was selected; for FUBAR, additional parameters under “advanced options” were set to their default values.

Residues under diversifying positive selection were mapped on the AA MSA of MCR subvariants and on the MCR-1 protein 3D structural model constructed based on the *Neisseria meningitidis* lipooligosaccharide phosphoethanolamine transferase EptA, using the Phyre2 server (accessed June 20, 2021) (Kelley et al., 2015; Anandan et al., 2017). The AA and the encoding nucleotide sequences of the residues predicted to have evolved under positive selection in different MCR subvariants were used to construct sequence logos. Briefly, the NT_btn_-MSA and the encoded AA-based MSA of the 98 known *mcr* subvariants were constructed using MUSCLE (Edgar, 2004) with default settings in Geneious version 2019.2.3 (Biomatters, Auckland, New Zealand). Aligned sequences of the 19 AA residues and codons (57 bases) identified by MEME were extracted from the corresponding MSA across the 98 *mcr* subvariants in Geneious and saved as separate MSA FASTA files. MSA FASTA files were used to construct AA and nucleotide graphical representation logos via the WebLogo3 server (accessed May 26, 2022) (Crooks et al., 2004).

### 2.9 *In silico* structural modeling

Structural modeling and structure editing were done as we have previously described (Carroll et al., 2019). Briefly, structural modeling of *mcr-1* was done using the Phyre2 server (accessed June 20, 2021) (Kelley et al., 2015) based on *Neisseria meningitidis* lipooligosaccharide phosphoethanolamine transferase EptA (Anandan et al., 2017). The protein structure was viewed and annotated using UCSF ChimeraX (Pettersen et al., 2021). The ESPript 3 server (version 3.0.8) was used to align secondary structure information onto the MSA (Robert and Gouet, 2014).

### 2.10 Collection of phenotypic colistin resistance data

Colistin minimum inhibitory concentration (MIC) data were collected from the literature as follows (Supplementary Table S8) (Di Pilato et al., 2016; Liu et al., 2016; Xavier et al., 2016; AbuOun et al., 2017; Borowiak et al., 2017; Carattoli et al., 2017; Ling et al., 2017; Liu et al., 2017; Lu et al., 2017; Poirel et al., 2017; Tijet et al., 2017; Yang et al., 2017; Yin et al., 2017; Zhao et al., 2017; Alba et al., 2018; Carattoli et al., 2018; Chavda et al., 2018; Dortet et al., 2018; Duggett et al., 2018; Eichhorn et al., 2018; Fernandes et al., 2018; Garcia-Graells et al., 2018; Hammerl et al., 2018; Kieffer et al., 2018; Li et al., 2018; Liu et al., 2018; Poirel et al., 2018; Rebelo et al., 2018; Shen et al., 2018; Teo et al., 2018; Wang et al., 2018b; Wang et al., 2018c; Wise et al., 2018; Xiang et al., 2018; Xu et al., 2018b; Xu et al., 2018c; Yang et al., 2018; Abdul Momin et al., 2019; Bitar et al., 2019; Chavda et al., 2019; Cui et al., 2019; Deshpande et al., 2019; Hadjadj et al., 2019; Li et al., 2019; Long et al., 2019; Wang et al., 2019; Yang et al., 2019; Yuan et al., 2019; Zhang et al., 2019; Zheng et al., 2019; Bitar et al., 2020; Cha et al., 2020; Fan et al., 2020; Garcia-Menino et al., 2020; Hatrongjit et al., 2020; Lei et al., 2020; Martins-Sorenson et al., 2020; Neumann et al., 2020; Ngbede et al., 2020; Wang et al., 2020; Anyanwu et al., 2021; Hu et al., 2021; Khanawapee et al., 2021; Leangapichart et al., 2021; Snyman et al., 2021; Stosic et al., 2021; Uddin et al., 2021; Yu et al., 2021). A search of NCBI’s PubMed database (https://pubmed.ncbi.nlm.nih.gov/) was conducted on August 13, 2021, using keywords [*mcr* and colistin resistance], and the search results were exported to an EndNote library (EndNote 20, Clarivate, Philadelphia, USA). A similar search was performed on the Web of Science Core Collection database (Clarivate, Philadelphia, USA), and the search results were used to amend the EndNote library. Duplicate entries in the EndNote library were deleted, and full-text PDF files were downloaded using EndNote “Find Full Text”. When needed, supplementary materials were downloaded and incorporated into the PDF file.

The EndNote library was searched for each *mcr* subvariant using the subvariant designation (e.g., *mcr-3.3*) with the “Any Field + PDF with Notes” search criteria. The search results were sorted by date to identify the primary publication(s) describing the subvariant for the first time. The full-text articles were visually scanned to identify the publication(s) describing (i) the identification of the subvariant and (ii) the phenotypic characterization of colistin resistance. The “Materials and Methods” and “Results” sections that described colistin resistance phenotypic data were examined carefully, and colistin MIC data were collected. We opted to use MIC values determined by the broth microdilution method, which is recommended by the European Committee on Antimicrobial Susceptibility Testing (EUCAST) (Jayol et al., 2018). We excluded MIC data determined by agar dilution and gradient agar disk diffusion (colistin gradient strip zone of inhibition) methods, which have been suggested to be less accurate in determining colistin resistance levels when compared to the broth microdilution method (Chew et al., 2017; Garcia-Menino et al., 2020; Yusuf et al., 2020). If no data were identified in the EndNote library for specific *mcr* subvariants, we performed an additional search using Google Scholar, using the given subvariant’s designation.

### 2.11 Colistin MIC ancestral state reconstruction

A ML phylogeny was constructed via IQ-TREE version 1.6.10 (Nguyen et al., 2015), using: (i) a NT_btn_-MSA as input (see section “Construction of maximum likelihood phylogeny of *mcr* and *mcr*-like genes” above), with *mcr* subvariants that lacked reliable native colistin MICs in the literature excluded (see section “Collection of phenotypic colistin resistance data” above); (ii) the best-fitting nucleotide substitution model selected using ModelFinder (i.e., based on Bayesian information critera [BIC] values, the TIM3e+G4 model) (Yang, 1994); (iii) 1,000 replicates of the ultrafast bootstrap approximation (Minh et al., 2013). Native MIC values associated with *mcr* subvariants, which were reported in the literature (denoted here as *MIC*_*Native*_; see Supplementary Table S8 and section “Collection of phenotypic colistin resistance data” above) were transformed using the following, where log refers to the natural logarithm:

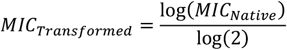

The ML phylogeny was supplied as input to the fastAnc function in the phytools version 0.7.80 R package (Revell, 2012), and the associated *MIC*_*Transformed*_ values were treated as a continuous character for which ancestral states were estimated (with “vars” and “CI” set to “True” so that variances and 95% confidence intervals would be computed for ancestral state estimates, respectively). The contMap function in phytools was used to plot the ML phylogeny with the mapped continuous character.

## 3 Results

### 3.1 *mcr*-like genes are distributed across a wide range of Gram-negative bacterial taxa

A protein BLAST (BLASTP)-based search of *mcr*-like genes in NCBI’s IPG database showed that *mcr*-like and phosphoethanolamine transferase (PET)-like genes were distributed across a wide range of Gram-negative bacterial taxa (Figure 1). More than 69,000 BLASTP hits against known MCR subvariants were detected in whole-genome sequencing (WGS) data spanning 256 bacterial genera (via the mOTUs taxonomy), including *Escherichia, Shigella, Salmonella, Klebsiella, Aeromonas, Moraxella, Enterobacter, Xanthomonas*, and *Pseudomonas* (Supplementary Table S2). Among the 69,814 total BLASTP hits to *mcr*-like genes identified here, 15,321 (21.9%) corresponded to genes annotated as *eptA* (i.e., the associated protein was annotated in NCBI with the term “EptA” or “eptA”; Supplementary Table S2).

**Figure 1.**
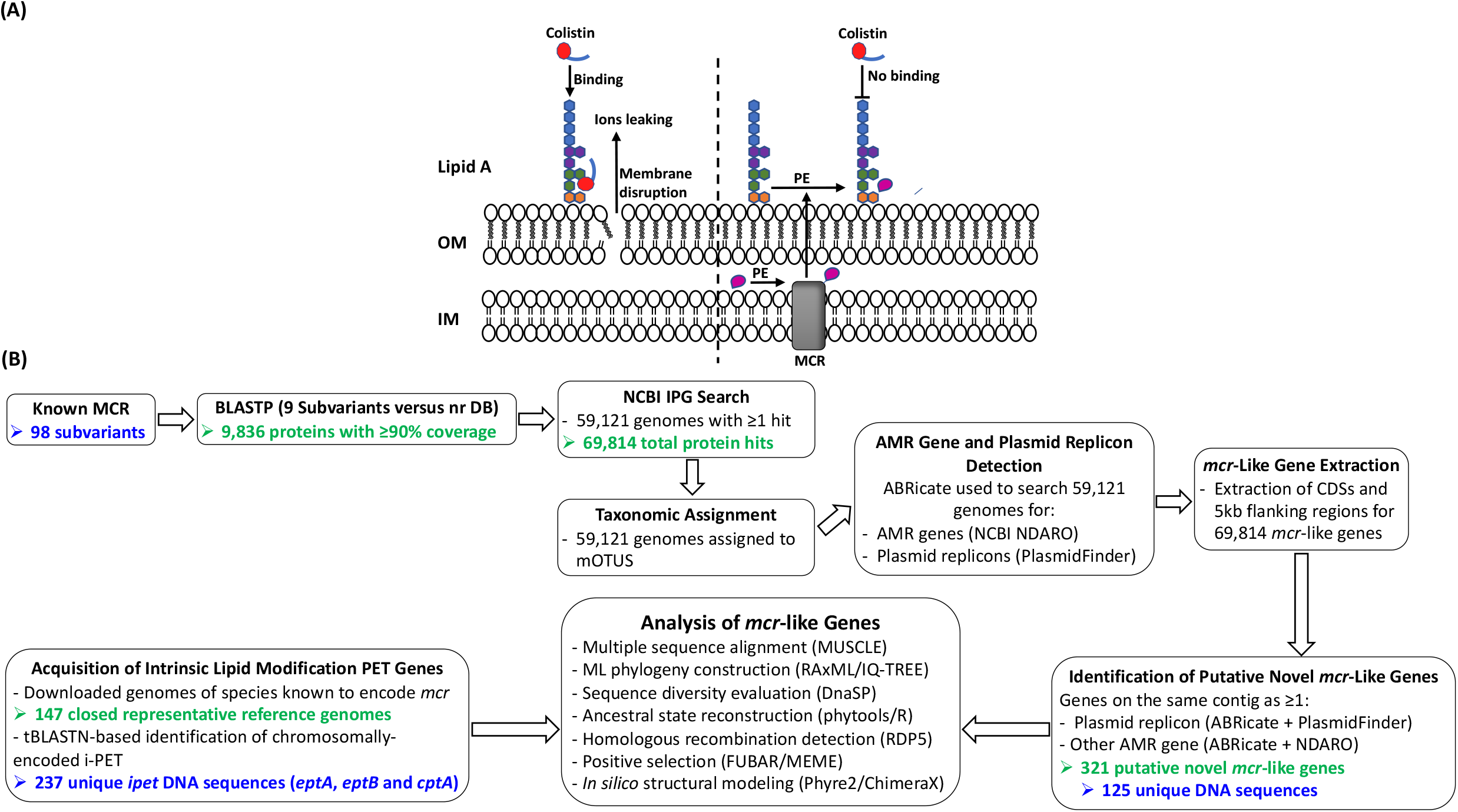
(A) Schematic diagram of colistin’s mode of action and resistance mechanism. Colistin, a cationic antimicrobial peptide, binds to the negatively charged lipid A, displacing membrane-bound cations and disrupting membrane integrity by inserting the hydrophobic tail into the membrane’s lipid (panel A, left). Lipid A modification neutralizes the negative membrane charge, thus reducing colistin binding and cell susceptibility to colistin (panel A, right). Abbreviations: IM, inner membrane; MCR, mobilized colistin resistance protein; OM, outer membrane; PE, phosphoethanolamine. (B) Schematic outline of the main methodologies used to acquire and characterize *mcr* and *mcr*-like genes from publicly available whole-genome sequencing (WGS) data (see the Materials and Methods section for details). The number of *mcr* genes, putative novel *mcr*-like genes, and/or intrinsic lipid modification phosphoethanolamine transferase (PET)-encoding genes (*ipet*) produced at relevant steps are shown in green text; the number of such genes used in final analyses are shown in blue text. Abbreviations: AMR, antimicrobial resistance; BLASTP, protein basic local alignment search tool; CDSs, coding sequences; DB, database; IPG, Identical Protein Group; MCR, mobilized colistin resistance amino acid sequences; ML, maximum likelihood; mOTUs, marker gene-based operational taxonomic units; NCBI, National Center for Biotechnology Information; NDARO, National Database of Antibiotic Resistant Organisms; nr, non-redundant protein sequence; (i)-PET, (intrinsic) phosphoethanolamine transferases; tBLASTN, translated nucleotide basic local alignment search tool.

*mcr* genes are typically plasmid-borne, while intrinsic lipid modification PET-encoding genes (*ipet*) are typically chromosomally encoded; thus, we hypothesized that *mcr*-like genes detected on the same contig as a plasmid-like origin of replication and other antimicrobial resistance (AMR) genes were more likely to encode PET associated with colistin resistance. Initially, we identified 321 BLASTP hits to *mcr*-like genes, which met these critera (defined here as “putative novel *mcr*-like genes”; Figure 1B and Supplementary Table S3). Sequence similarities showed that the 321 putative novel *mcr*-like genes represented 125 unique nucleotide sequences, which were selected for further analyses (Figure 1B and Supplementary Tables S3 and S7).

Within a back-translated nucleotide multiple sequence alignment (NT_btn_-MSA) of 98 known *mcr* subvariants, plus the 125 unique, putative novel *mcr-*like genes identified here (Supplementary Tables S3 and S6), an average pairwise nucleotide diversity per site (π) of 0.42675 was observed, with an average number of pairwise nucleotide differences per sequence (κ) of 623.052 over 1,776 sites. The relatively high π value observed here is not surprising, as it has been shown previously that *mcr* subvariants encompass an extensive degree of genetic diversity (Khedher et al., 2020); this is also evident within a NT_btn_-MSA constructed using only known *mcr* subvariants (i.e., π = 0.3614 and κ = 553.669 over 1,776 sites; Supplementary Table S6).

A maximum likelihood (ML) phylogeny inferred from the NT_btn_-MSA of *mcr* and putative novel *mcr*-like genes showed a robust separation of clades and subclades representing different *mcr* families (Figure 2 and Supplementary Figure S1). Specifically, *mcr* subvariants clustered into two distinct phylogenetic clades: (i) Clade A included subvariants belonging to the *mcr-1, mcr-2, mcr-5*, and *mcr-6* families, and (ii) Clade B included all remaining *mcr* subvariants (Figure 2 and Supplementary Figure S1). Interestingly, both clades contained multiple putative novel *mcr*-like genes (Figure 2 and Supplementary Figure S1). While some putative novel *mcr*-like genes were closely related to known *mcr* subvariants, several putative novel *mcr*-like genes formed phylogenetic clades that were distinct from known *mcr* subvariants (Figure 2 and Supplementary Figure S1).

**Figure 2.**
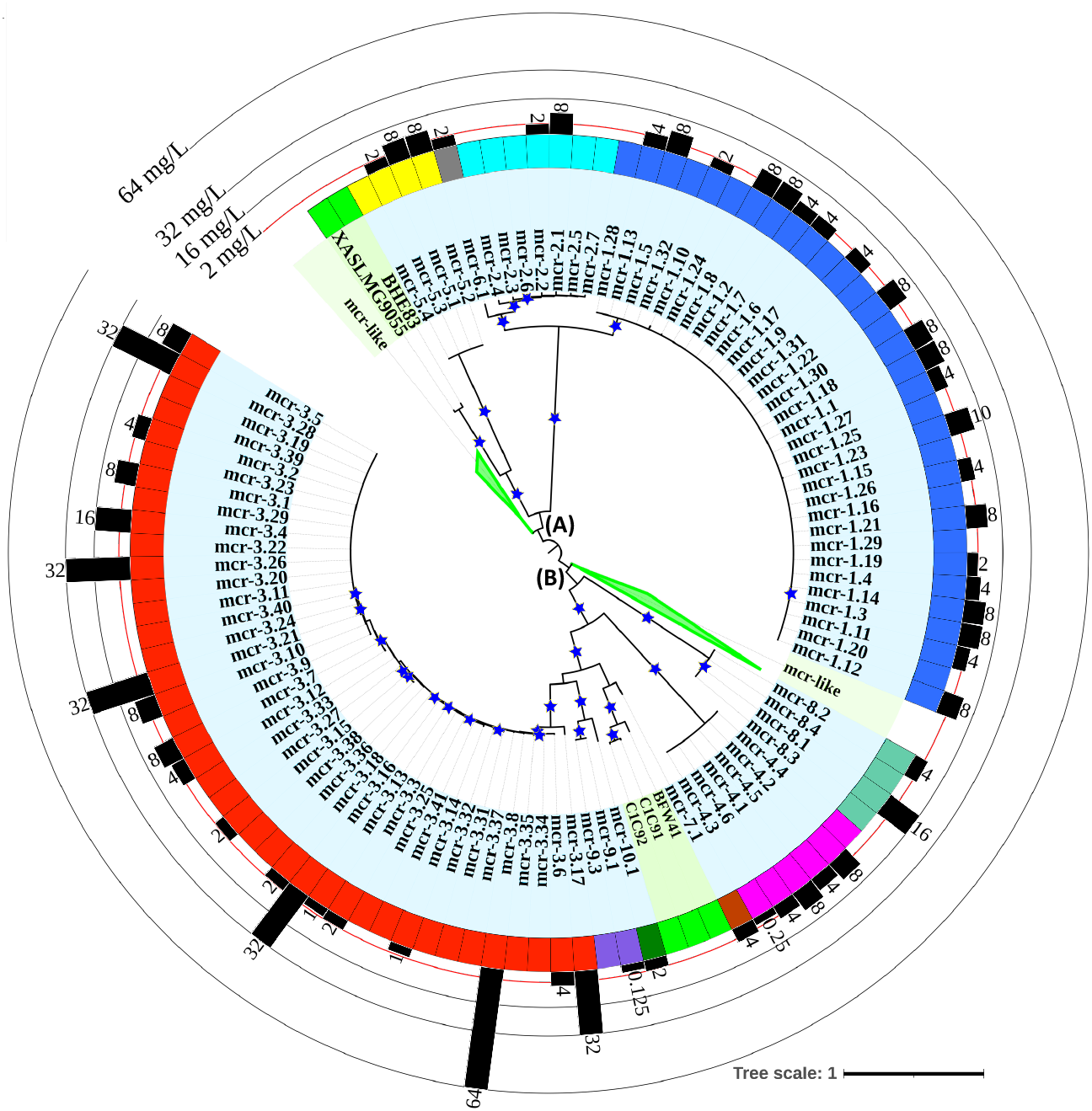
Maximum likelihood (ML) phylogeny inferred from a nucleotide back-translation-based multiple sequence alignment (NT_btn_-MSA) of (i) 98 *mcr* subvariants and (ii) 125 unique sequences of *mcr*-like genes encoded on the same contig as ≥1 plasmid origin of replication and ≥1 other antimicrobial resistance (AMR) gene (i.e., “putative novel *mcr*-like genes”). Sequences were aligned using MUSCLE. The ML phylogeny was constructed with RAxML, using the GTRGAMMA nucleotide substation model and 100 bootstrap replicates. The tree was edited using the iTOL web server (https://itol.embl.de/) and rooted at the midpoint, with branch lengths reported in substitutions per site. Branches with bootstrap values ≥70% are denoted by blue stars. Clades exclusively composed of putative novel *mcr*-like genes were collapsed for clarity (green branches; see Supplementary Figure S1 for a fully expanded tree). Tip label shading corresponds to known *mcr* subvariants (light blue) and putative novel *mcr*-like genes identified in this study (light green). The colors of the inner ring represent different *mcr* families (*mcr-1* to *-10*). The outer graph represents the maximum reported colistin minimum inhibitory concentration (MIC) values of native strains encoding different *mcr* subvariants. The colistin breakpoint established by the Clinical and Laboratory Standards Institute (CLSI) is 2 mg/L (red line). Isolates with colistin MIC ≥2 mg/L are considered colistin-resistant. Metadata associated with each gene can be found in Supplementary Tables S3 and S6; colistin MIC values can be found in Supplementary Table S8.

### 3.2 Some *mcr* subvariants are more closely related to *eptA* than to other *mcr* subvariants

Given the wide range of genetic diversity observed among *mcr* subvariants, amino acid (AA) sequence similarities among MCR subvariants also varied widely as expected, with similarities ranging from 59.3% to 100% and identities ranging from 29.7% to 99.8% (Supplementary Tables S9 and S10). However, to gain further insight into MCR diversity within the context of intrinsic lipid modification PET (i-PET), we identified and aggreagated 237 chromosomally encoded i-PET (i.e., EptA, EptB, and CptA; Supplementary Table S5). The i-PET identified here were derived from 147 genomes of species known to encode *mcr* (e.g., *Aeromonas, Citrobacter, Cronobacter, Enterobacter, Escherichia, Klebsiella, Moraxella, Proteus*, and *Salmonella* spp.; Supplementary Tables S4 and S5) (Nang et al., 2019). The nucleotide and AA sequences of the 237 chromosomally encoded i-PET identified here were compared to those of the 98 known MCR subvariants, as well as the sequences of the 125 putative novel MCR-like genes identified here (*n* = 460 total PET; Supplementary Table S7). Based on nucleotide sequences, all 460 PET queried here displayed extensive sequence diversity (i.e., π = 0.50571 and κ = 450.085; Supplementary Table S7).

Based on AA sequences, MCR subvariants shared a relatively low degree of AA similarity with EptB and CptA, ranging from 46% to 58% and 42% to 49%, respectively (Supplementary Tables S9 and S10). Notably, however, MCR subvariants differed in terms of the degree of AA similarity they shared with EptA: MCR AA similarities to EptA ranged from 61% to 76%, and some MCR subvariants shared a higher degree of AA similarity to EptA than to other MCR subvariants (Supplementary Tables S9 and S10). MCR-1.1, for example, showed average similarities of 64.5 ±1.5% to EptA and 76.1±18.1% to other MCR subvariants (Supplementary Tables S9 and S10). MCR-8.1, on the other hand, showed average similarities of 72.8 ± 1.1% to EptA and 70 ±7% to other MCR subvariants (Supplementary Tables S9 and S10). Thus, it is evident that sequence similarity cannot sufficiently discriminate between canonic EptA and MCR.

The topology of a ML phylogeny produced using nucleotide sequences of *mcr*, putative novel *mcr*-like genes, and *ipet* (*n* = 460 total PET-encoding nucleotide sequences; Supplementary Table S7) provided further evidence that some *mcr* subvariants**—**including putative novel *mcr*-like subvariants identified here**—**were more closely related to *eptA* than to other *mcr* subvariants (Figure 3 and Supplementary Figure S2). Overall, 26 of 125 putative novel *mcr-*like genes (20.8%) clustered among known *mcr* genes in the ML phylgeny (Figure 3 and Supplementary Figure S2). Among these, two and three genes were found to be most closely related to *mcr-5* and *mcr-7* (groups c and f, Figure 3), respectively; the remaining 21 putative novel *mcr-*like genes formed two distinct phylogenetic groups (groups d and e, Figure 3). Notably, however, a large proportion of putative novel *mcr-*like genes (99 out of 125, 79.2%), identified as such based on similarity to known *mcr* genes and predicted localization on a plasmid, were more closely related to *eptA* than to known *mcr* subvariants (Figure 3 and Supplementary Figure S2). Thus, additional criteria are likely needed to differentiate bona fide *mcr* from *eptA*.

**Figure 3.**
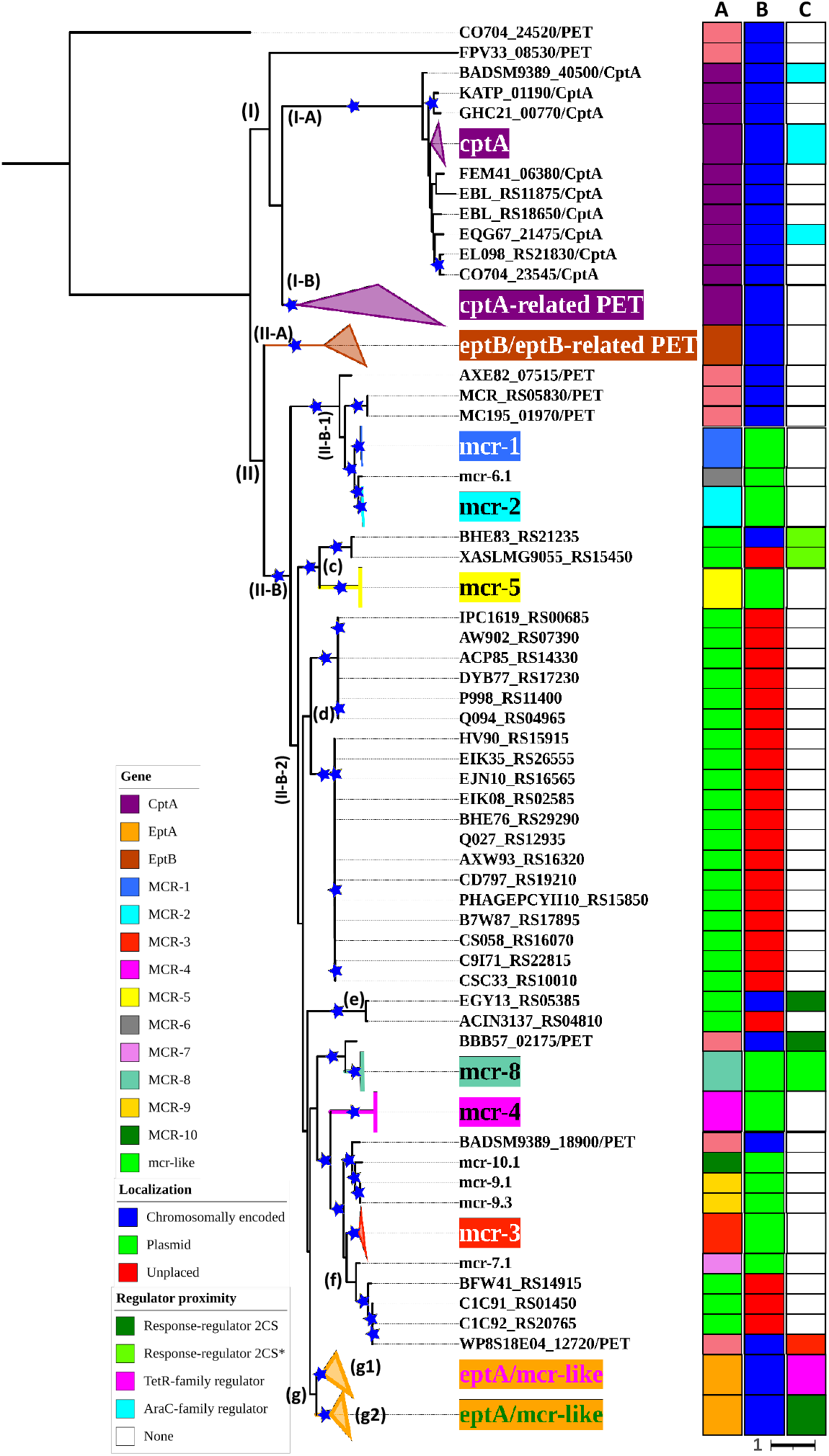
Maximum likelihood (ML) phylogeny inferred from a nucleotide back-translation-based multiple sequence alignment (NT_btn_-MSA) of (i) 98 *mcr* subvariants, (ii) 125 unique sequences of *mcr*-like genes encoded on the same contig as ≥1 plasmid origin of replication and ≥1 other antimicrobial resistance (AMR) gene (i.e., “putative novel *mcr*-like genes”), and (iii) 237 chromosomally encoded phosphoethanolamine transferase (*ipet*) genes. Sequences were aligned using MUSCLE. The ML phylogeny was constructed with RAxML, using the GTRGAMMA nucleotide substation model and 100 bootstrap replicates. The tree was edited using the iTOL web server (https://itol.embl.de/) and rooted at the midpoint, with branch lengths reported in substitutions per site. Branches with bootstrap values ≥70% are denoted by blue stars. Clades exclusively composed of genes from the same family were collapsed and color-coded as shown in the left color legend key (see Supplementary Figure S2 for a fully expanded tree). Color-coded regions in strip (A) denote *mcr* families and *eptA, eptB*, and *cptA* homologues. Color-coded regions in strip (B) denote gene localization as chromosomally encoded (blue), plasmid-encoded (green), or unplaced (red). Color-coded regions in strip (C) represent the regulatory system juxtaposing phosphoethanolamine transferase (PET)-encoding genes: dark green regions represent genes encoded in the same operon with a two-component response sensor-regulatory system; light green regions represent genes encoded divergent of a two-component response sensor-regulatory system; magenta regions represent genes encoded adjacent to a TetR-type regulator; cyan regions represent genes encoded adjacent to an AraC-type regulator; white regions represent genes encoded as a single gene operon and no upstream or downstream regulatory protein.

### 3.3 Colistin resistance and intrinsic lipid modification PET are genetically and functionally diversified

Within the ML phylogeny of *mcr*, putative novel *mcr*-like genes, and *ipet* (*n* = 98, 125, and 237 genes, respectively; Supplementary Table S7), PET-encoding genes clustered into two superclades, including (i) a superclade with 105 taxa, which included *cptA* genes (designated [I] in Figure 3), and (ii) a superclade with 354 taxa, which included *eptA, eptB, mcr* and putative novel *mcr*-like genes (designated [II] in Figure 3). As previously shown (Harper et al., 2017), *eptA, eptB*, and *cptA* genes clustered into distinct phylogenetic clades that correlate with the enzymes’ acceptor substrate specifities (Figure 3 and Supplementary Figure S2). Interestingly, a large subset of chromosomally encoded *ipet* formed a distinct subclade (designated [I-B] in Figure 3) within the *cptA*-containing superclade, which may indicate that genes in subclade [I-B] encode enzymes with acceptor substrate specificities similar to, but not identical to, the substrate specificity of CptA.

Genes in superclade [II] formed two distinct phylogenetic groups: (i) an *eptB*-containing clade [II-A], and (ii) a *mcr*, putative novel *mcr*-like, and *eptA*-containing clade [II-B] (Figure 3). Genes in clade [II-B] further clustered into two subclades, where subvariants belonging to the *mcr-1, mcr-2*, and *mcr-6* families clustered in subclade [II-B-1], while subvariants belonging to the remaining *mcr* families, as well as *eptA*, clustered in subclade [II-B-2] (Figure 3).

While most *mcr* are plasmid-encoded, chromosomally-encoded *mcr* genes have been reported (Ling et al., 2017; Dortet et al., 2018; Stogios et al., 2018; Wang et al., 2021). Interestingly, several chromosomally encoded PET genes from *Moraxella* spp. (loci numbers AXE82_07515, MC195_01970 and MCR_RS05830), *Kosakonia sacchari* (locus number BBB57_02175), *Aeromonas caviae* (locus number WP8S18E04_12720), and *Buttiauxella agrestis* (locus number BADSM9389_18900) clustered with known *mcr* families and thus are likely to encode MCR-related functions (Figure 3 and Supplementary Figure S2).

### 3.4 The genomic contexts of *mcr* and *eptA* are widely diverse

To identify other criteria that may aid in identifying *mcr* genes, we studied the genomic context of *mcr, eptA, eptB, cptA*, and the putative novel *mcr*-like genes identified in this study (Figure 3, Supplementary Figure S2, and Supplementary Table S11). We found that all 59 *eptA* and 99 *eptA*-related putative novel *mcr-*like genes identified here clustered into two phylogenetic groups, where (i) the “g1” group included 37 genes encoded adjacent to a putative TetR/AcrR family transcriptional regulator, and (ii) the “g2” group included 121 genes encoded in an operon with a two-component sensor histidine kinase-response regulator system (Figure 3, Supplementary Figure S2, and Supplementary Table S11).

In comparison, a gene encoding an AraC-type transcriptional regulator was frequently encoded 70-200 nucleotides downstream of the *cptA* coding region (25 of 33 queried genes, 75.8%; Figure 3, Supplementary Figure S2, and Supplementary Table S11). *eptB*, on the other hand, was solely encoded as single gene operons (based on 48 genes queried here). Interestingly, all 107 *cptA*-related genes in clade [I-B] were encoded as single gene operons (Figure 3, Supplementary Figure S2, and Supplementary Table S11).

In contrast to i-PET-encoding genes, the genomic context of *mcr* was highly variable among different *mcr* families (Figure 3, Supplementary Figure S2, and Supplementary Table S11). *mcr-1* subvariants, for example, were encoded in a putative operon upstream of a PAP2 family lipid A phosphatase-encoding gene, while *mcr-3* subvariants were encoded upstream of diacylglycerol kinase-encoding gene *dgkA*. Interestingly, *mcr-7* was located 125 nucleotides upstream of a *pap2-dgkA* operon. Recently, it was shown that *pap*2 and *dgkA* genes encoding a PAP2 family lipid A phosphatase and diacylglycerol kinase, respectively, play a role in *mcr-*dependent colistin resistance through recycling and modification of lipid metabolism and byproducts (Choi et al., 2020; Gallardo et al., 2020; Purcell et al., 2022). Only *mcr-8* subvariants were encoded adjacent to, but divergent from, an operon encoding a two-component sensor histidine kinase-response regulator. Comparatively, subvariants belong to the remaining *mcr* families were encoded as single gene operons (Figure 3, Supplementary Figure S2, and Supplementary Table S11).

### 3.5 Structurally and functionally important disulfide bonds are differentially conserved among different MCR and i-PET families

Sequence analysis of MCR, EptA, EptB, and CptA proteins showed that five disulfide bonds, which were identified in the EptA structure (Anandan et al., 2017), were differentially conserved among different MCR families (Supplementary Figure S2 and Supplementary Table S7). Previously, structural and mutational analyses of the *E. coli* EptC, a CptA homolog, revealed that the inability of EptC to confer colistin resistance was due to the lack of several disulfide bonds (Zhao et al., 2019). Zhao et al. (Zhao et al., 2019) suggested that the disulfide bonds function synergistically to restrain the flexibility of different loops, especially a loop located near the active site, which forms part of a potential substrate entry tunnel. Sequence analysis showed that CptA homologs lacked all predicted disulfide bonds, while EptA and EptB homologs were predicted to contain five and three disulfide bonds, respectively (Supplementary Figure S2 and Supplementary Table S7). Interestingly, only three disulfide bonds were conserved in MCR-1, MCR-2, and MCR-6 subvariants, while five disulfide bonds were conserved in subvariants belonging to the other MCR families (Supplementary Table S7). However, the role that disulfide bond number variability plays in the functional diversity of MCR subvariants remains unknown.

### 3.6 *mcr* subvariants confer varying levels of colistin resistance

It has been shown that colistin resistance levels vary among *mcr*-encoding strains, and colistin minimum inhibitory concentration (MIC) data are lacking for multiple *mcr* subvariants in the NCBI Reference Gene Catalog (see Nang, et al. for a recent extensive literature survey of colistin MIC values associated with different bacterial isolates) (Nang et al., 2019). Indeed, we found that native strains encoding different *mcr* subvariants have markedly diverse colistin MIC levels (Figures 2 and 4 and Supplementary Table S8). For example, colistin MIC values of 2, 4, 8, and 10 mg/L were reported for isolates expressing *mcr-1* subvariants (Supplementary Table S8). Further, colistin MIC values of 2, 4, 8, 16, 32, and 64 mg/L were reported for *mcr-3* harboring strains, despite the relatively high genetic similarity of *mcr-3* subvariants (i.e., physiologically characterized *mcr-3* subvariants share 88.7% to 99.8% identity and 97.6% to 100% similarity on the AA level; Figures 2 and 4 and Supplementary Table S8). Moreover, colistin MIC values varied among species expressing the same *mcr* subvariant (Figure 4 and Supplementary Table S8). *E. coli* and *Salmonella* spp. encoding the *mcr-2.1* subvariant, for example, had colistin MIC values of 8 and 4 mg/L, respectively (Xavier et al., 2016; Garcia-Graells et al., 2018; Poirel et al., 2018). Interestingly, an *E. coli* wild strain encoding the *mcr-3.10* subvariant on a plasmid had a colistin MIC value of 8 mg/L, while *Aeromonas caviae* and *Proteus mirabilis* encoding a chromosomal *mcr-3.10* subvariant had a colistin MIC of 32 mg/L (Wang et al., 2018c).

**Figure 4.**
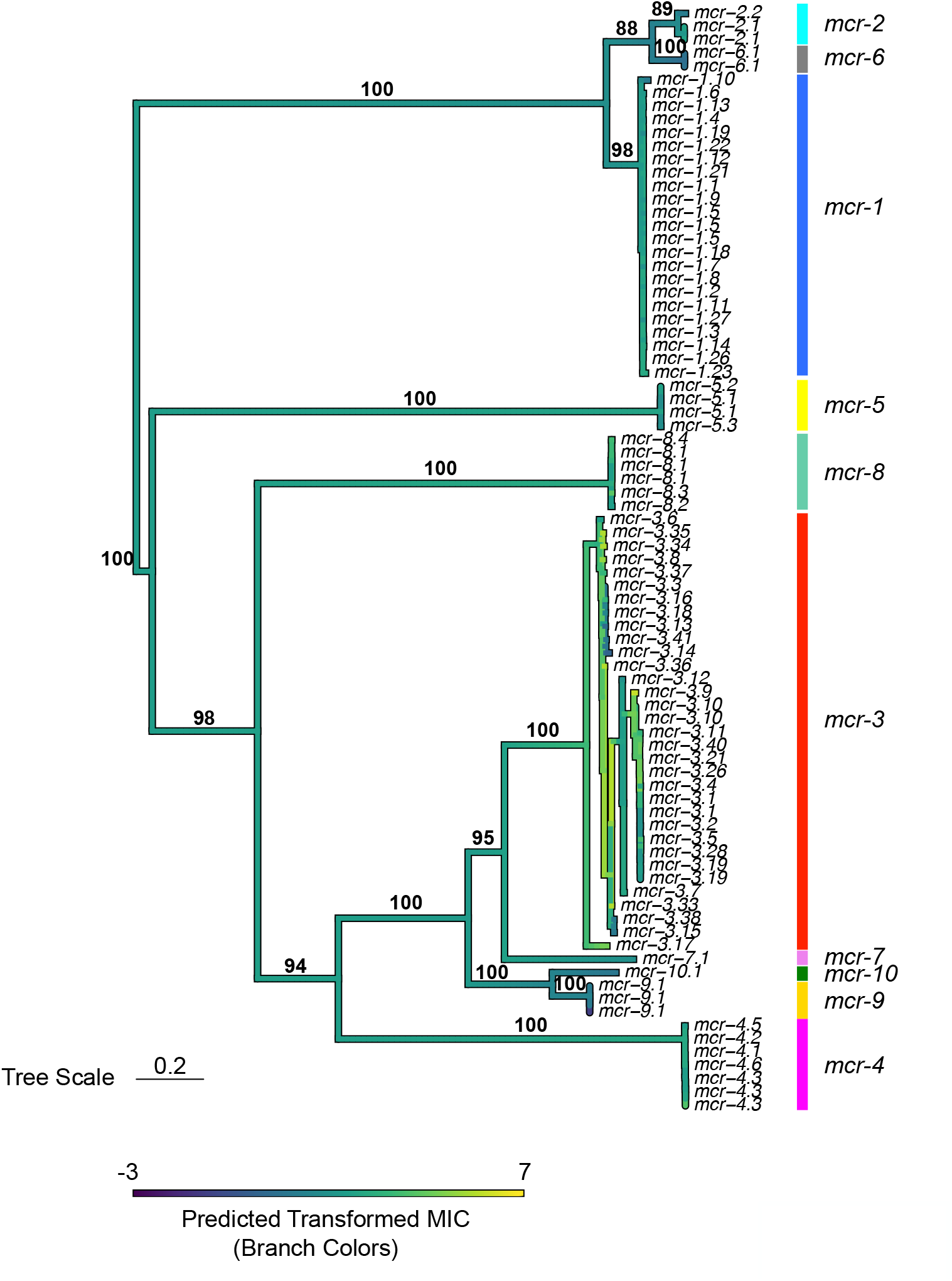
Ancestral state reconstruction results based on colistin minimum inhibitory concentration (MIC) values obtained for *mcr* subvariants in their native strains. MIC values were obtained from the literature and transformed by taking the natural logarithm of the native values and dividing by the natural logarithm of 2 (i.e., transformed MIC values; see the Materials and Methods section for details). Branch colors denote predicted transformed MIC values, which were estimated using the fastAnc function in the phytools package in R. Predicted transformed MIC values were mapped to a maximum likelihood (ML) phylogeny of the selected MCR subvariants using the contMap functrion in the phytools R package. The ML phylogeny was constructed using IQ-TREE, with a nucleotide back-translation-based multiple sequence alignment (NT_btn_-MSA) of 83 *mcr* subvariants supplied as input (*mcr* subvariants that did not have a reliable colistin MIC in a native strain reported in the literature were excluded; Supplementary Table S8). The phylogeny is rooted along the midpoint, with branch lengths reported in substitions per site. Clade labels to the right of the phylogeny denote *mcr* families (*mcr-1* to *-10*). Branch lables (boldfaced black text) denote branch support percentages obtained using the ultrafast bootstrap approximation (selected for readability).

Notably, multiple *mcr* subvariants have been characterized by heterologous expression in standard laboratory strains (Supplementary Table S8). We observed extensive variability in colistin MIC values among *E. coli* laboratory strains heterologously expressing different *mcr* subvariants (Supplementary Table S8). In heterologous expression systems, several factors are likely to affect AMR levels, including the strains’ genetic background, plasmid copy number, level and regulation of gene expression, and cellular toxicity resulting from protein overexpression (Wagner et al., 2008; Schlegel et al., 2012; Tietgen et al., 2018).

### 3.7 Recombination plays a limited role in the evolution of *mcr*

Thirteen potential recombination events were initially identified in a NT_btn_-MSA of *mcr*, putative novel *mcr*-like, and *ipet* genes via multiple methods in RDP5 (*n* = 460 total PET-encoding genes; Supplementary Figure S3 and Supplementary Table S7). Specifically, initial evidence for recombination among *mcr-3* subvariants was identified (i.e., events 1-3; Supplementary Figure S3). However, visual inspection of the recombination events indicated that recombinant fragments in these three events shared similar breakpoints, indicating that these fragments represented a single recombination event (Supplementary Figure S3). Moreover, within the *mcr* and *ipet* phylogeny, *mcr-3* subvariants identified in recombination events 1-4 clustered closely into subclades with relatively short terminal branches (Figure 3 and Supplementary Figure S3). Similar observations were found for subvariants associated with recombination events 5-8 and 10-13 (Figure 3 and Supplementary Figure S3), indicating that the recombination signals identified in events 1-8 and 10-13 (i.e., all events except for event 9) might have been caused by evolutionary processes other than homologous recombination, such as inter-lineage and inter-site mutation-rate variation (Bertrand et al., 2016; Martin et al., 2021).

Comparatively, recombination event 9, which included *mcr-7, mcr-3.34*, and a newly identified putative novel *mcr*-like gene from *Aeromonas caviae* (gene locus ID C1C91_01450 and NCBI Nucleotide accession CP025706.1), was identified by six different recombination detection methods in RDP5 (pairwise homoplasy index [PHI] raw *P* = 0.035; Supplementary Figure S3). The PET phylogenies (Figures 2 and 3) revealed that the putative recombinant and parents were divergent and had relatively long terminal branches, suggesting that recombination event 9 may have played a role in the evolution of some *mcr-3* and *mcr-7* subvariants (Supplementary Figure S3). Overall, these results posit a limited role for homologous recombination in the evolution and diversification of *mcr* subvariants; however, it is feasible that some recombination events (e.g., between closely related sequences or genes not included in the analysis) were involved in the evolution of *mcr*.

### 3.8 Positive selection likely contributed to the evolution of specific amino acid residues that may induce localized structural variation among MCR subvariants

The role of positive selection in the evolution of genes can be assessed by estimating the non-synonymous to synonymous substitution rate (*dN*/*dS*) ratio (ω), where an overall ω > 1 indicates positive selection. Alternatively, ω = 1 or ω < 1 suggests neutral or negative selection, respectively (Murrell et al., 2012). Overall, an average ω value of 0.198 across the 98 known *mcr* subvariants was observed (via DnaSP; Supplementary Table S6), suggesting that *mcr* evolved under negative selection. Indeed, FUBAR (the Fast, Unconstrained Bayesian AppRoximation method, which infers *dN*/*dS* on a per-site basis) (Murrell et al., 2013) predicted that 212 AA sites may have evolved under negative selection (FUBAR posterior probability ≥ 0.99; Supplementary Figure S4 and Supplementary Table S12). To predict a possible structural role for the AA sites that evolved under diversifying selection, we mapped these AA residues on a MCR-1 to -10 MSA supplemented with MCR-1 secondary structure elements (Supplementary Figure S4). We found that AA sites that may have evolved under negative diversifying selection were distributed among the three structural components of MCR-1. Specifically, 36% of the AA residues of the membrane-anchored domain, 18% of the AA residues of the bridging region, and 44% of the AA residues of the catalytic domain were predicted to evolve under diversifying negative selection (Supplementary Figure S4). Using FUBAR, no sites were predicted to evolve under positive selection at posterior probability ≥ 0.99.

While an overall *dN*/*dS* ratio < 1 indicates that positive selection did not likely contribute to the overall evolution of *mcr*, positive selection may have still played a role in the evolution of subvariant-specific and branch-specific AA sites. Indeed, relatively few genes with *dN*/*dS* ratio > 1 are expected to exist (Murrell et al., 2012). Hence, we used a mixed-effect model of evolution (MEME) approach to identify episodic diversifying positive selection at the level of the individual subvariants, a branch, or a set of branches. MEME showed that positive selection may have played a role in the diversification of 19 subvariant-specific AA sites within 34 branch-specific nodes distributed among two partitions in the phylogeny of 98 known *mcr* subvariants (MEME raw *P* < 0.05; Figure 5 and Supplementary Table S13). Among the 19 AA sites, 11 residues were located within the membrane-anchored domain, one in the bridging helix, one in the interdomain flexible loop, and six in the catalytic domain (Figure 5). Interestingly, MEME identified multiple events in the *mcr-2* and *mcr-9* family phylogeny branches (Figure 5).

**Figure 5.**
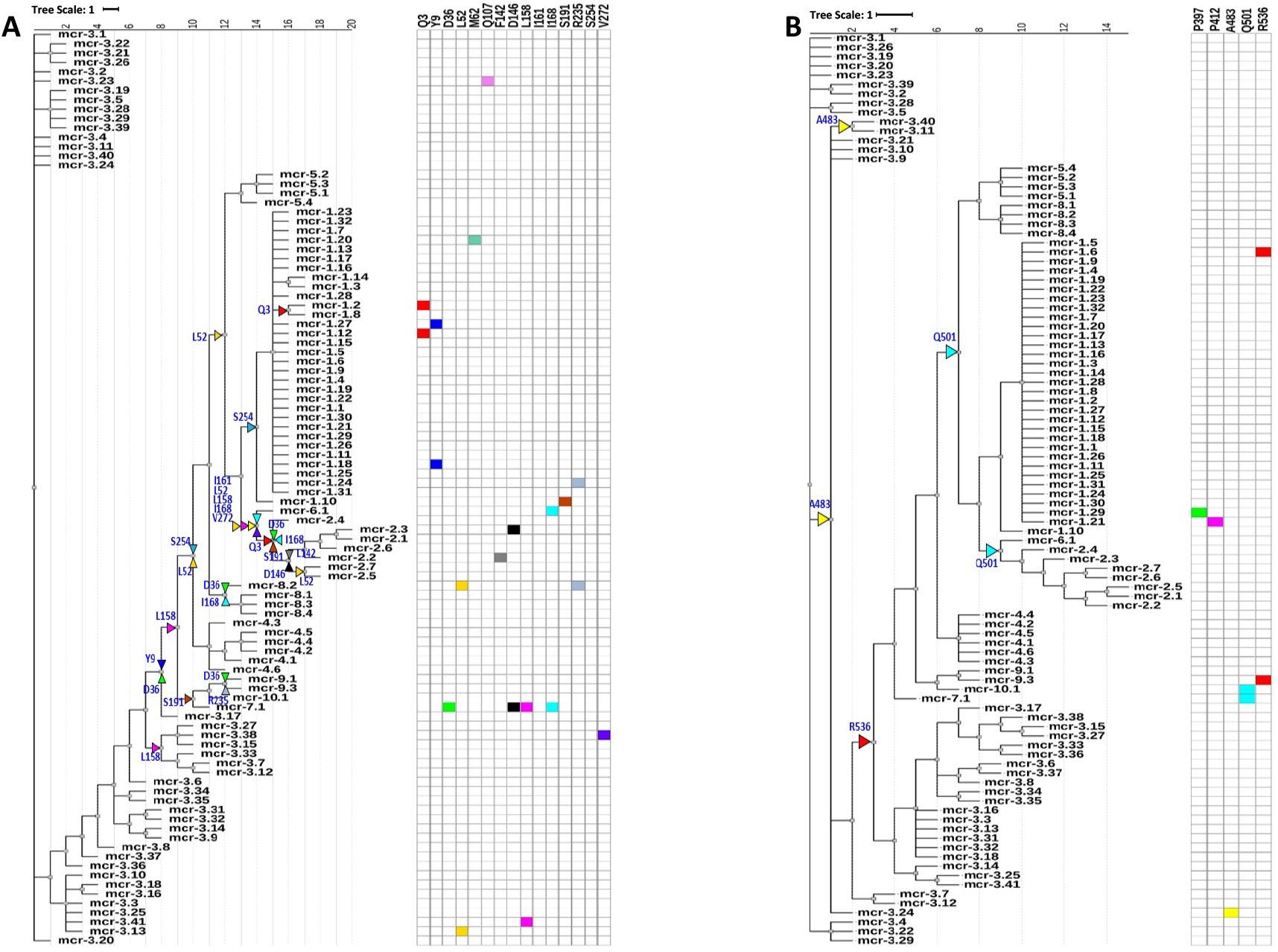
Identification of amino acid (AA) residues under branch-specific positive selection. A phylogeny of 98 known *mcr* subvariants produced and partitioned by GARD (https://www.datamonkey.org/gard) shows partition 1 (A) and partition 2 (B) branch-specific AA sites evolving under positive selection. Sequences were aligned using MUSCLE, and the resulting multiple sequence alignment (MSA) was supplied as input to GARD. The partitioned dataset produced by GARD was supplied as input to MEME (mixed-effect model of evolution-based diversifying selection analysis; https://www.datamonkey.org/meme), which was used to identify AA residues under branch-specific positive selection. The tree (default output/rooting produced by MEME, with branch lengths reported in substitions per site) was edited using the iTOL web server (https://itol.embl.de/). Color-coded regions and triangles represent AA residues under subvariant-specific and branch-specific positive selection, respectively.

While MEME did not identify positive selection evidence for AA residues involved in zinc and pEtN binding, the analysis predicted that several structurally critical residues evolved under diversifying positive selection (Supplementary Table S13). Subsequently, we mapped these residues on the structural model of MCR-1 to understand the structural-functional relationship of AA diversity at these loci (Figures 6A and 6B). We found that several of the AA residues that evolved under diversifying selection were located in structurally important regions, including (i) Gln^107^, which was located at a periplasmic loop connecting the third and fourth transmembrane segments juxtaposing the substrate entry tunnel, (ii) Ser^191^ and Arg^235^, which were located at a bridging region that connects the N-terminal membrane domain and the C-terminal catalytic domain, and (iii) Pro^397^, Pro^412^, and Ala^483^, which were located adjacent to the active site and substrate entry tunnel, respectively (Figures 6A and 6B). Subsequently, we evaluated the sequence diversity of the codons associated with these 19 AA residues among *mcr* subvariants (Figure 6C). Overall, we found that 47 sites out of 57 nucleotides (i.e., 19 codons) were polymorphic among all *mcr* families. In contrast, coding sequences of the 19 AA residues were highly conserved among different subvariants of each *mcr* family (e.g., π values of synonymous and non-synonymous substitutions were 0.47189 and 0.348 among all subvariants, 0.00276 and 0.01728 among *mcr-1* subvariants, and 0.02769 and 0.02938 among *mcr-3*).

**Figure 6.**
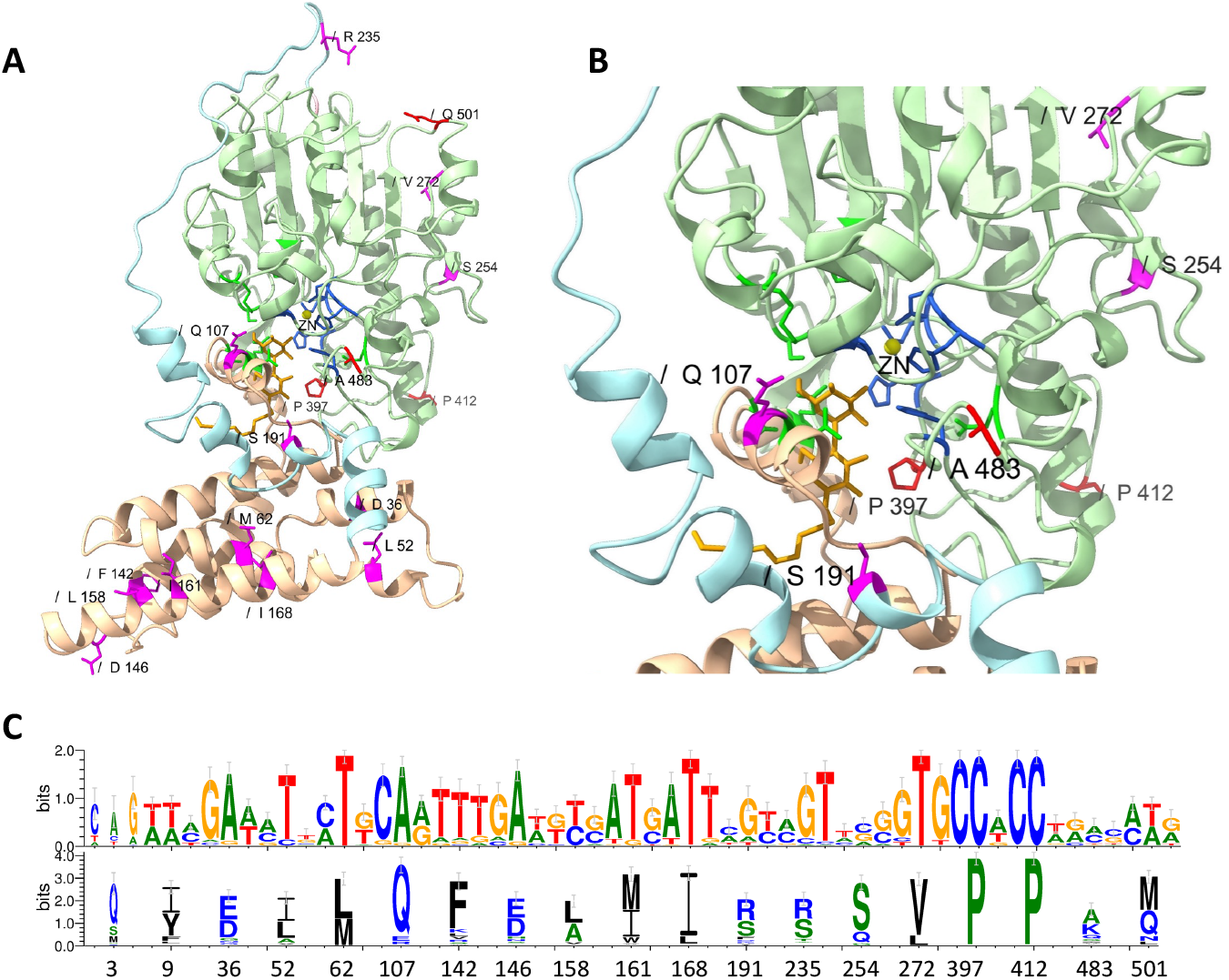
Structural localization and conservation of amino acid (AA) residues under branch-specific positive selection. (A) Structural model of the MCR-1 protein, constructed based on *Neisseria meningitidis* phosphoethanolamine transferase EptA (Anandan et al., 2017). The MCR structural model was constructed using the Phyre2 server, and the structure was viewed and edited using UCSF ChimeraX. The structural model shows the transmembrane-anchored domain (yellow) and the soluble periplasmic catalytic domain (light green) connected by a bridging helix and extended loop (light blue). AA residues involved in zinc (yellow circle) and phosphoethanolamine (pEtN) binding are colored dark blue and green, respectively. Branch-specific AA residues evolving under positive selection are shown in magenta (partition 1) and red (partition 2). (B) Close-up of the MCR active site showing the localization of branch-specific AA residues under positive selection in relation to the active site and the substrate entry tunnel. The structure shows Gln^107^, which is located at a periplasmic loop connecting the third and fourth transmembrane segments juxtaposing the substrate entry tunnel; Ser^191^, which is located at a bridging region that connects the N-terminal membrane domain and the C-terminal catalytic domain; and Pro^397^, which is located adjacent to the active site. (C) Web-logo of nucleotide and AA sequences showing the conservation of 19 AA residues identified by MEME (https://www.datamonkey.org/meme) to have evolved under diversifying positive selection among the 98 known MCR subvariants.

### 3.9 Novel *mcr* subvariants and families may be present among publicly available bacterial genomes

Using analyses that considered sequence similarity, genomic localization, and genomic context, we identified multiple putative novel *mcr-*like genes among publicly available bacterial genomes, which likely encode novel *mcr* subvariants or families (Supplementary Table S3). For example, the *mcr*-like genes of *Xanthomonas campestris* (locus tag BHE83_RS21235) and *Xanthomonas phaseoli* (locus tag XASLMG9055_RS15450) are likely to be *mcr-5* subvariants. This is corroborated by the fact that these genes are phylogenetically close to other *mcr-5* subvariants and are localized adjacent to a PAP2 family lipid A phosphatase-encoding gene. Additionally, *mcr*-like genes from *Aeromonas hydrophila* and *Aeromonas caviae* (loci tags BFW41_RS14915, C1C91_RS01450, and C1C92_RS20765) are likely to encode novel *mcr-7* subvariants. Comparatively, eighteen *mcr*-like genes from *Pseudomonas aeruginosa* (group d, Figure 3) and two *mcr*-like genes from *Acinetobacter baumannii* and *Acinetobacter* spp. FDAARGOS 493 (group e, Figure 3) are likely to encode novel *mcr* families. In addition to sharing a relatively low degree of sequence similarity with known *mcr* families, all of these genes lacked the regulatory elements that we found to be universally identified within the genomic context of *eptA*. Thus, experimental phenotypic analyses are needed to define the potential roles of these genes in colistin resistance and assess their phenotypic diversity.

## 4 Discussion

Overall, our data show that (i) PET-encoding genes can be detected across a wide range of Gram-negative bacterial species and are genetically and functionally diverse; (ii) the evolution and diversification of *mcr* is likely the result of a multifaceted process that includes positive selection, gene mobilization, and diversification of regulatory pathways and gene contexts; (iii) a holistic approach that considers genomic context, in addition to sequence similarity and genomic localization, may aid in the identification of novel *mcr* families and subvariants.

### 4.1 Genetically and functionally diverse PET-encoding genes can be detected across a wide range of Gram-negative bacterial species

Previous studies (Nang et al., 2019; El-Sayed Ahmed et al., 2020; Khedher et al., 2020) have shown that PET-encoding genes are distributed across a range of Gram-negative bacteria; here, we identified >69,000 genes homologous to colistin resistance-conferring PET in >1,000 bacterial species (based on the mOTUs taxonomy; Supplementary Table S2). Using a subset of PET-encoding genes composed of (i) all 98 known MCR-encoding genes, (ii) 125 genes encoding putative novel MCR-like proteins, and (iii) 237 chromosomal i-PET-encoding genes, we provide further insight into the genetic diversity of colistin resistance-conferring and instrinsic PET alike (Figure 3). Like previous studies (Carroll et al., 2019; Khedher et al., 2020), we have shown that MCR subvariants are genetically diverse. However, here we observed that some *mcr* subvariants are more closely related to chromsonal i-PET-encoding *eptA* than to some other *mcr* subvariants (Figure 3). Previous studies have hypothesized that *mcr* evolved from *eptA* through mobilization of an *eptA* gene copy flanked by transposable insertion elements (Kieffer et al., 2017; Wang et al., 2018a). While the exact origin of the *mcr* ancestor is still unknown, *Moraxella* spp. have been suggested as a potential source (Kieffer et al., 2017).

In addition to being genetically diverse, PET-encoding genes are functionally diverse as well. Specifically, different *mcr* subvariants can confer a wide range of colistin resistance levels (Nang et al., 2019). Using previously published colistin MIC values associated with native *mcr*-encoding strains, we found that the genetic diversity of *mcr* does not strictly correlate with colistin resistance phenotypic diversity. For example, colistin resistance levels are markedly diverse among strains encoding different *mcr-3* subvariants, while colistin resistance levels are less diverse among strains encoding *mcr-1* subvariants, despite comparable intra-family sequence similarity levels (Supplementary Table S8). While colistin resistance phenotypic heterogeneity conferred by different *mcr* subvariants is evident, these data should be cautiously viewed. Colistin’s primary mode of action is the displacement of LPS-stabilizing cations followed by insertion into the outer membrane and cell disruption; however, it has been suggested that oxidative damage plays a role in colistin-dependent cell killing (Trimble et al., 2016; El-Sayed Ahmed et al., 2020; Cianciulli Sesso et al., 2021). As such, the highly diverse and species-specific oxidative stress response (Touati, 2000; Imlay, 2003; O’Connor and McClean, 2017) can alter the susceptibility of bacterial cells to colistin treatment. Thus, a clear understanding of the extent to which *mcr* genetic diversity contributes to colistin resistance heterogeneity remains a major challenge.

### 4.2 Evolution and diversification of *mcr* is the result of a multifaceted process

Our data suggest that multiple factors might be involved in the evolution and functional diversification of *mcr*, including the diversification of the genomic context and the regulatory control of gene expression. We found that all tested chromosomally encoded *eptA* and putative novel *mcr*-like genes most closely related to *eptA* were localized adjacent to an AraC-type regulator or a two-component sensor histidine kinase-response regulator system. With the exception of *mcr-8*, no specific regulator was found to be associated with the remaining *mcr* subvariants. In contrast, we found that lipid A phosphatase-encoding *pap2* and diacylglycerol kinase-encoding *dgkA* were frequently encoded adjacent to many *mcr* subvariants. Specifically, *mcr-1* and *mcr-2* subvariants were encoded adjacent to *pap2*, while *mcr-3* and *mcr-8* subvariants were encoded adjacent to *dgkA*. Recently, a detailed mutational analysis indicated that *pap2* and *dgkA* play a significant role in *mcr*-dependent colistin resistance, possibly through recycling and modification of lipid metabolism and byproducts (Choi et al., 2020; Gallardo et al., 2020; Purcell et al., 2022). Moreover, it was shown that a transposon insertional element has horizontally transferred the *mcr-1*-*pap2* region into different plasmid backbones (Li et al., 2016; AbuOun et al., 2017; Zhang et al., 2019). The role of *pap2* in *mcr* function is further corroborated by the recent identification of a *Sutterella wadsworthensis* gene encoding a single polypeptide with an N-terminal Pap2 domain and C-terminal MCR-like domain (Andrade et al., 2021). Hence, it is evident that the genomic contexts of *eptA* and *mcr* are differentially associated with the presence of regulatory protein or accessory enzymes of lipid metabolism. *eptA* are exclusively encoded adjacent to transcription regulators, while *mcr* are frequently encoded adjacent to enzymes for the recycling and modification of lipid metabolism and byproducts.

Additionally, our data suggest that positive selection likely contributed to the diversification of specific *mcr* subvariants, as we found that diversifying positive selection might have played a role in the evolution of several AA residues adjacent to functionally and structurally important protein regions. While AA residues predicted to evolve under positive selection are not part of the active site and substrate-binding site, sequence variations at these loci are likely to affect the enzyme function, such as alteration of substrate accessibility, specificity, and enzyme efficacy. For example, we predicted that branch-specific positive selection might be involved in the diversification of Ser^191^ and Arg^235^ in the interdomain-connecting region. Recently, it has been shown that the conformational flexibility of EptA is essential in enzyme-substrate recognition (Anandan et al., 2021). The conformational changes of EptA are governed by a highly conserved domain structure that offers extensive flexibility between the membrane-bound and the periplasmic catalytic domains (Anandan et al., 2017; Anandan and Vrielink, 2020). Moreover, Anandan et al. (Anandan et al., 2021) proposed that the bridging helix acts as a hinge region that enables extensive conformational changes between the two domains, “opening up” the catalytic domain to allow access to the considerably large lipid A substrate. Hence, diversification of Ser^191^ and Arg^235^ is likely to influence the overall conformational flexibility of the enzyme. For example, at position 191, the relatively small polar Ser may offer better conformational flexibility compared to other AA residues identified by MEME at the same locus, such as the non-polar aromatic Phe or Trp, and the negatively charged Asp or Glu (Supplementary Table S13).

Similarly, diversification of Gln^107^, Pro^397^, Pro^412,^ and Ala^483^, which are located in several periplasmic loops juxtapose and adjacent to the substrate entry tunnel and the active site (Anandan et al., 2017; Anandan and Vrielink, 2020), is likely to influence accessibility to the substrate entry tunnel, which may affect enzyme activity. Overall, diversification of these AA residues might contribute to the phenotypic diversity of MCR subvariants in colistin resistance. However, detailed site-directed mutational analyses will be required to define the role of positive selection in phenotypic diversity of MCR.

Overall, our data raise the intriguing possibility that *eptA* can give rise to colistin resistance genes through a multifaceted process that includes genetic evolution and diversification of genomic context, mobilization, and regulatory pathways. However, the relative contribution of these factors in the evolution of *mcr* remains unknown.

### 4.3 A holistic approach that considers sequence, structure, genomic localization, and genomic context may aid in the discovery of novel *mcr* families and subvariants

In addition to providing insight into the genetic and functional diversity of *mcr* and other PET, we identified 125 unique, putative novel *mcr*-like genes, which were located on the same contig as (i) ≥1 plasmid replicon and (ii) ≥1 additional AMR gene (Supplementary Table S3). It may be tempting to speculate on the colistin resistance-conferring potential of the putative novel *mcr*-like genes identified here; however, the discrimination between colistin resistance-conferring *mcr* and intrinsic lipid modification *eptA* remains a clear challenge, as evident from (i) the high genetic diversity among *mcr* subvariants and (ii) the absence of phenotypic data associated with a large portion of the *mcr* phylogeny. Furthermore, it is essential to note the limitations of the criteria used to select putative novel *mcr*-like genes in this study. Specifically, it is possible that a plasmid replicon could be detected in a chromosomal contig. Previous studies have shown that 10% of bacterial genomes do not contain a single circular chromosome, and instead, each genome is composed of several essential replicons, including secondary chromosomes and chromids, which use a plasmid-like origin of replication for propagation (Harrison et al., 2010; diCenzo and Finan, 2017). In these cases, the approach used here could incorrectly classify such a contig as plasmid-associated, should it contain a replicon and an additional AMR gene. Thus, the putative novel *mcr*-like genes identified here may encode i-PET enzymes, or they may constitute distinct bona fide groups of colistin resistance-conferring *mcr* genes. Regardless, functional analyses will be essential to assign biological functions to these genes.

Overall, inferring AMR phenotypes from bacterial WGS data remains challenging (Tyson et al., 2015; Ransom et al., 2020). Functional discrimination between *eptA* and *mcr* would optimally require physiological characterization, which may involve the construction of gene-specific mutants or molecular cloning in a heterologous expression system (Gao et al., 2016; Liu et al., 2016; Carroll et al., 2019). However, these approaches are time-consuming and require a hard-to-achieve standardization process to ensure similar gene expression and active protein levels. Moreover, it has been shown that heterologous overexpression of *P. aeruginosa eptA* (Cervoni et al., 2021) or dysregulation of *S*. Typhimurium *eptA* transcription (Vaara et al., 1981; Olaitan et al., 2014) lead to colistin resistance. Hence, the identification of other bioinformatic criteria that can be used to discriminate between *mcr* and *eptA* can improve the ability to detect colistin resistance genes in bacterial WGS data and provide an important basis for selecting genes for phenotypic characterization. In the future, we imagine that holistic approaoches that consider multiple criteria (e.g., genomic context, in addition to sequence and structural homology to known *mcr* subvariants, localization on mobile genetic elements) will aid researchers in the identification and prioritization of novel *mcr* families and subvariants from bacterial WGS data.

## 5 Author Contributions

AG and LMC performed all computational analyses. AG, MW, and LMC designed the study and co-wrote the manuscript.

## 7 Data Availability Statement

NCBI accession numbers for all genes and genomes included in this study are available in Supplementary Tables S1-S7.

