## Supplementary Figure S1 for "More than *mcr*: Canonical Plasmid- and Transposon-Encoded Mobilized Colistin Resistance (*mcr*) Genes Represent a Subset of Phosphoethanolamine Transferases"

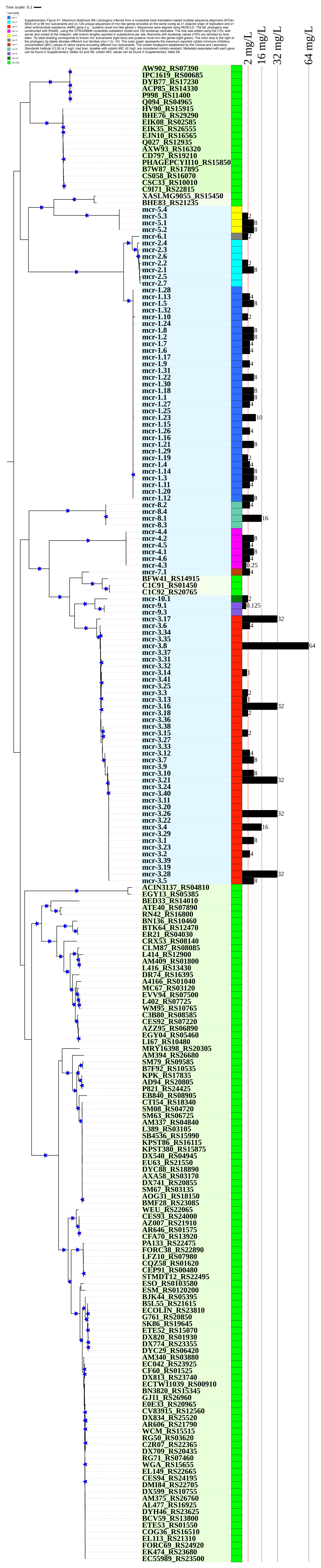


Supplementary Figure S1. Maximum likelihood (ML) phylogeny inferred from a nucleotide back-translation-based multiple sequence alignment (NTbtn-MSA) of (i) 98 mcr subvariants and (ii) 125 unique sequences of mcr-like genes encoded on the same contig as ≥1 plasmid origin of replication and ≥1 other antimicrobial resistance (AMR) gene (i.e., "putative novel mcr-like genes"). Sequences were aligned using MUSCLE. The ML phylogeny was constructed with RAxML, using the GTRGAMMA nucleotide substation model and 100 bootstrap replicates. The tree was edited using the iTOL web server and rooted at the midpoint, with branch lengths reported in substitutions per site. Branches with bootstrap values ≥70% are denoted by blue stars. Tip label shading corresponds to known mcr subvariants (light blue) and putative novel mcr-like genes (light green). The color strip to the right of the phylogeny tip labels denotes different mcr families (mcr-1 to -10). The outer graph represents the maximum reported colistin minimum inhibitory concentration (MIC) values of native strains encoding different mcr subvariants. The colistin breakpoint established by the Clinical and Laboratory Standards Institute (CLSI) is 2 mg/L (red line). Isolates with colistin MIC ≥2 mg/L are considered colistin-resistant. Metadata associated with each gene can be found in Supplementary Tables S3 and S6; colistin MIC values can be found in Supplementary Table S8.
