## Supplementary Figure S2 for "More than *mcr*: Canonical Plasmid- and Transposon-Encoded Mobilized Colistin Resistance (*mcr*) Genes Represent a Subset of Phosphoethanolamine Transferases"

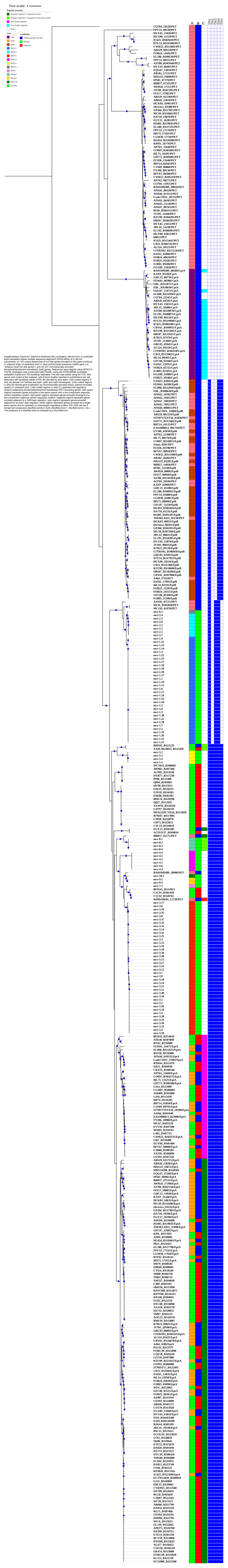

Supplementary Figure S2. Maximum likelihood (ML) phylogeny inferred from a nucleotide back-translation-based multiple sequence alignment (NTbtn-MSA) of (i) 98 mcr subvariants, (ii) 125 unique sequences of mcr-like genes encoded on the same contig as ≥1 plasmid origin of replication and ≥1 other antimicrobial resistance (AMR) gene (i.e., “putative novel mcr-like genes”), and (iii) 237 chromosomally encoded phosphoethanolamine transferase (ipet) genes. Sequences were aligned using MUSCLE. The ML phylogeny was constructed with RAxML, using the GTRGAMMA nucleotide substation model and 100 bootstrap replicates. The tree was edited using the iTOL web server and rooted at the midpoint, with branch lengths reported in substitutions per site. Branches with bootstrap values ≥70% are denoted by blue stars. Color-coded regions in strip (A) denote mcr families and eptA, eptB, and cptA homologues. Color-coded regions in strip (B) denote gene localization as chromosomally encoded (blue), plasmid-encoded (green) or unplaced (red). Color-coded regions in strip (C) represent the regulatory system juxtaposing phosphoethanolamine transferase (PET)-encoding genes: dark green regions represent genes encoded in the same operon with a two-component response sensor-regulatory system; light green regions represent genes encoded divergent of a two-component response sensor-regulatory system; magenta regions represent genes encoded adjacent to a TetR-type regulator; cyan regions represent genes encoded adjacent to an AraC-type regulator; white regions represent genes encoded as a single gene operon and no upstream or downstream regulatory protein. DS1-DS5 are the 5 conserved consecutive disulfide bonds in EptA (disulfide bond 1, disulfide bond 2, etc.). The presence of a disulfide bond is indicated by a blue-filled box.
