## Supplementary Figure S3 for "More than *mcr*: Canonical Plasmid- and Transposon-Encoded Mobilized Colistin Resistance (*mcr*) Genes Represent a Subset of Phosphoethanolamine Transferases"

Supplementary Figure S3. Genes with evidence of homologous recombination as detected by RDP5 and PHI analysis.

| Event # | Sequence | RDP | GENE-CONV | Boot- scan | Maxchi | Chim-aera | SiSscan | 3Seq | PHI p value | 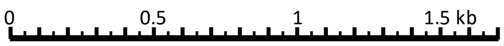 |
|---|---|---|---|---|---|---|---|---|---|---|
| 1-3[$] | *mcr*-3.33/*mcr*-3.9/*mcr*-3.10[R]<br>*mcr*-3.12/*mcr*-3.14/*mcr*-3.7[Mj]<br>*mcr*-3.3/ *mcr*-3.13/ *mcr*-3.16[Mn] | 1.03E-26 | 3.09E-23 | 2.44E-27 | 2.09E-13 | 9.61E-14 | 1.44E-21 | 4.62E-41 | <0.00001 | 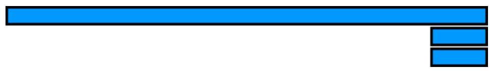 |
| 4[$] | *mcr*-3.1[R]<br>*mcr*-3.37[Mj]<br>*mcr*-3.17[Mn] | NS | 3.7E-18 | 4.9E-22 | 1.7E-11 | 1.1E-11 | 6.6E-16 | 6.4E-25 | <0.00001 | 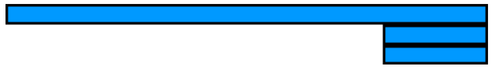 |
| 5 | PET_IC625_RS02610[R]<br>*yhbX*_D7U33_RS16100[Mj]<br>PET_HUX90_12110[Mn] | NS | 5.0E-09 | 6.5E-12 | 6.5E-11 | 1.4E-10 | 7.9E-16 | 3.3E-21 | 0.29 | 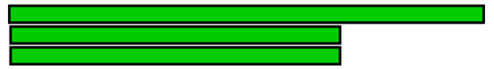 |
| 6[$] | *mcr*-like_DX820_RS01930[R]<br>*mcr*-like_ETE52_RS15070[Mj]<br>*eptA*_JRC41_19530[Mn] | NS | NS | NS | 2.8E-10 | 2.5E-04 | 1.7E-10 | 1.6E-13 | <0.00001 | 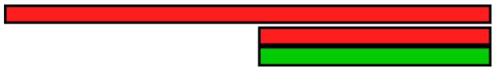 |
| 7[$] | *mcr*-like_SK86_RS19645[R]<br>*mcr*-like_AM340_RS03880[Mj]<br>*eptA*_JRC41_19530[Mn] | NS | NS | NS | 1.9E-03 | 1.9E-06 | 8.0E-10 | 1.2E-12 | <0.00001 | 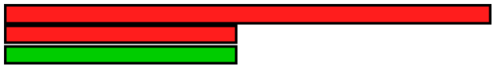 |
| 8[$] | *mcr*-like_P821_RS24425[R]<br>*eptA*_I8N75_17455[Mj]<br>*mcr*-like_EU63_RS21550[Mn] | 2.4E-06 | 8.5E-08 | 3.5E-07 | 4.4E-06 | 3.9E-08 | 3.7E-07 | 1.6E-11 | <0.00001 | 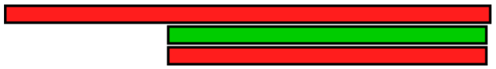 |
| 9 | *mcr*-7[R]<br>*mcr*-3.34[Mj]<br>*mcr*-like_C1C91_RS01450[Mn] | 9.25E-07 | NS | 9.09E-10 | 4.07E-12 | 7.16E-15 | 1.87E-30 | 2.55E-15 | **0.0348** | 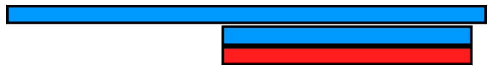 |
| 10[$] | *eptB*_FY206_01050[R]<br>*eptB*_electrica_00201[Mj]<br>PET_BFV67_00910[Mn] | NS | 3.9E-09 | 1.8E-05 | 4.8E-05 | 9.9E-04 | 1.0E-08 | NS | 3.9E-03 | 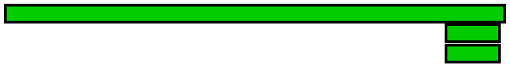 |
| 11 | *cptA*_HVX45_15055[R]<br>*cptA*_EL192_RS00875[Mj]<br>*cptA*_ D7U33_RS19990[Mn] | 4.7E-02 | NS | 3.8E-02 | 6.2E-04 | 2.3E-05 | NS | 4.9E-06 | 0.11 | 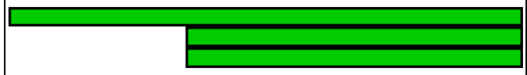 |
| 12[$] | *mcr*-like_EB840_RS08905[R]<br>*mcr*-like_EU63_RS21550[Mj]<br>*eptA*_I8N75_17455[Mn] | NS | 3.2E-03 | 2.2E-04 | 5.8E-05 | 4.3E-05 | 4.4E-07 | NS | <0.00001 | 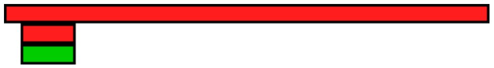 |
| 13 | *cptA*_JRC41_20400[R]<br>*cptA*_GBC03_05945[Mj]<br>*cptA*_ E4Z61_15935[Mn] | NS | NS | 2.7E-02 | 1.4E-03 | 4.4E-05 | 2.4E-04 | 9.8E-03 | 0.20 | 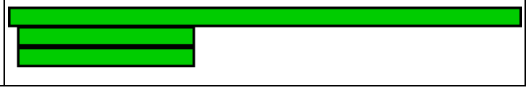 |

NS: No significant P-value was recorded for this recombination event using this method.
($) RDP analysis of this event indicates that recombination might not have caused this signal, while P-value indicates some evidence of recombination.
R: recombinant; Mj: major parent; Mn: minor parent
