## Supplementary Figure S4 for "More than *mcr*: Canonical Plasmid- and Transposon-Encoded Mobilized Colistin Resistance (*mcr*) Genes Represent a Subset of Phosphoethanolamine Transferases"

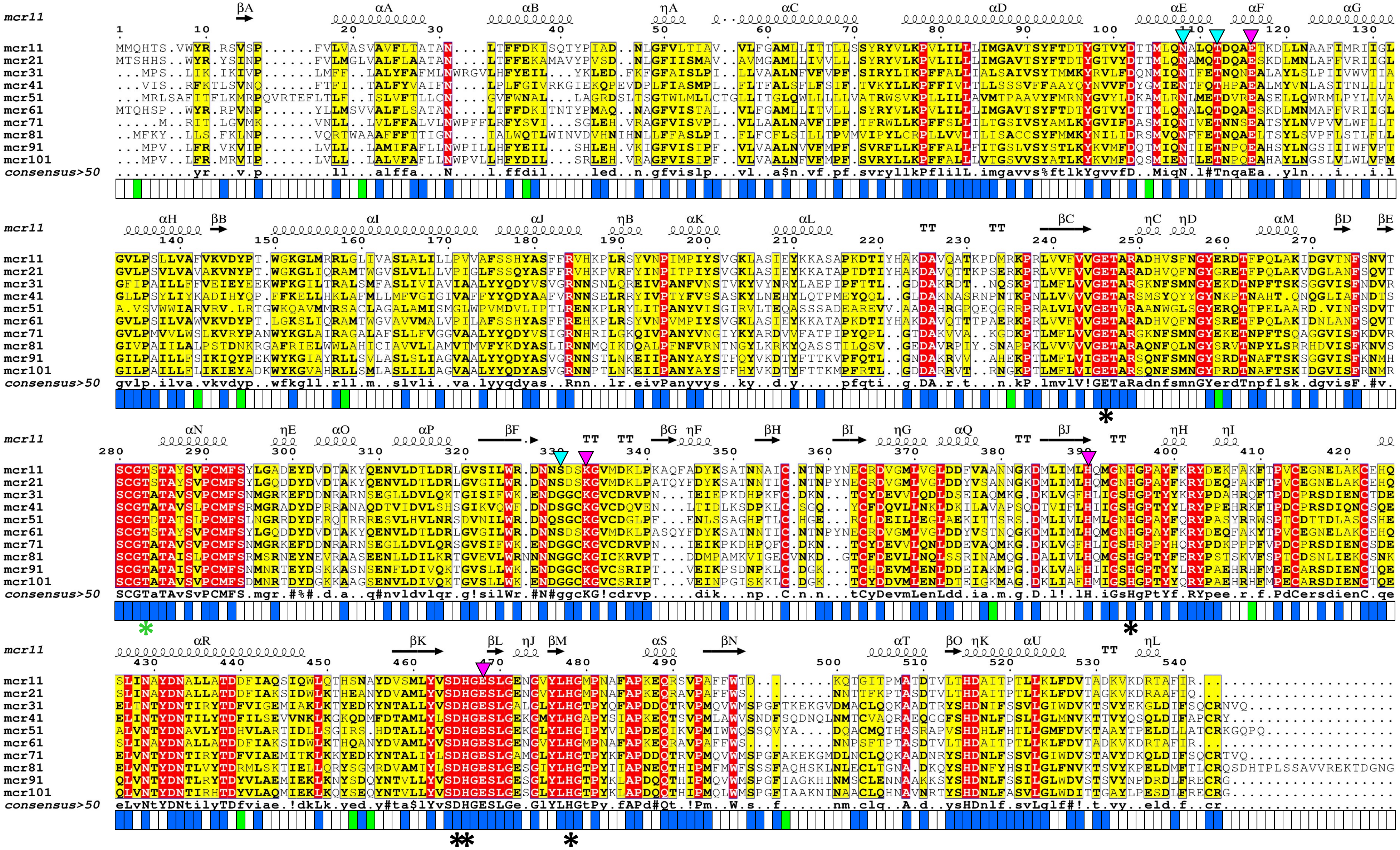

Supplementary Figure S4. Multiple sequence alignment (MSA) of MCR subvariants showing site-specific amino acid (AA) residues evolving under pervasive negative/purifying selection (at posterior probability values of > 0.99; blue boxes below MSA) and pervasive positive/diversifying selection (at posterior probability values of > 0.5; green boxes below MSA). The top track denotes MCR-1 secondary structure elements (alpha helixes and beta sheets) aligned to the MSA using the ESPript 3 server. Within the MSA, a strict identity (i.e., identical AA residue at a site) is denoted by a red box and a white character, while a yellow box around an AA residue denotes similarity across the groups. Conserved AA residues involved in zinc-binding are denoted by black stars below the MSA, while a green star identifies the catalytic threonine residue. AA residues involved in phosphoethanolamine (pEtN) binding are indicated by inverted triangles, where magenta and cyan triangles denote residues that are essential or partially essential for MCR function, respectively.
